# GxE PRS: Genotype-environment interaction in polygenic risk score models for quantitative and binary traits

**DOI:** 10.1101/2023.07.20.549816

**Authors:** Dovini Jayasinghe, Md. Moksedul Momin, Kerri Beckmann, Elina Hypponen, Beben Benyamin, S. Hong Lee

## Abstract

The use of polygenic risk score (PRS) models has transformed the field of genetics by enabling the prediction of complex traits and diseases based on an individual’s genetic profile. However, the impact of genotype-environment interaction (GxE) on the performance and applicability of PRS models remains a crucial aspect to be explored. Currently, existing GxE PRS models are often inappropriately used, which can result in inflated type 1 error rates and compromised results. In this study, we propose a novel GxE PRS model that correctly incorporates the GxE component to analyze complex traits and diseases. Through extensive simulations, we demonstrate that our proposed model outperforms existing models in terms of controlling type 1 error rates and enhancing statistical power. Furthermore, we apply the proposed model to real data, and report significant GxE effects. Specifically, we highlight the impact of our model on both quantitative and binary traits. For quantitative traits, we uncover the GxE modulation of genetic effects on body mass index (BMI) by alcohol intake frequency (ALC). In the case of binary traits, we identify the GxE modulation of genetic effects on hypertension (HYP) by waist-to-hip ratio (WHR). These findings underscore the importance of employing a robust model that effectively controls type 1 error rates, thus preventing the occurrence of spurious GxE signals. To facilitate the implementation of our approach, we have developed an innovative R software package called GxE PRS, specifically designed to detect and estimate GxE effects. Overall, our study highlights the importance of accurate GxE modeling and its implications for genetic risk prediction, while providing a practical tool to support further research in this area.

## Introduction

Human complex traits and diseases arise from a complex interplay between genetic and environmental factors. For example, conditions such as body mass index (BMI), cholesterol levels, diabetes, coronary artery disease (CAD), and hypertensive disorders are all influenced by both genetic and environmental factors, which can interact to modulate genetic effects on these outcomes. This concept is supported by evidence from twin and family-based studies ^1,2,3,4^.

In the genomic era, genome-wide association studies (GWAS) have made significant progress in identifying genetic variants associated with complex traits and diseases ^5^. Leveraging the summary statistics from GWAS, polygenic risk scores (PRSs) can be generated as a single measure that reflects an individual’s risk for a specific disease. PRSs are widely utilized in genomic prediction, providing valuable insights into disease susceptibility ^6^. Additionally, genome-wide environment interaction studies (GWEIS) have been employed to identify genetic variants whose effects are influenced by environmental factors ^7,8,9^. Incorporating gene-environment interactions (GxE) into genomic prediction, such as GxE PRS models, holds promise for improving the accuracy of predicting complex traits and diseases, particularly when modifiable environmental variables are available.

Existing methods for GxE PRS models face several challenges that need to be addressed. The current literature presents various GxE PRS models ^10,11,12,13^, where PRS term considered in the interaction component is, primarily constructed based on additive genetic effects only. However, this approach can substantially reduce the power to detect GxE in the target dataset, an issue that has received limited investigation ^14,15^. Another challenge in GxE PRS models is the potential for model misspecification ^16,17,18^. When higher-order polynomial effects of the environment or unknown non-genetic effects modulated by the environment are present, spurious GxE signals may arise. Unfortunately, this problem has not been extensively studied, highlighting the need for further investigation and understanding.

To tackle these challenges, we conducted extensive simulations to assess the potential improper application of existing GxE PRS models in terms of the type 1 error rate and statistical power. We also evaluated the impact of model misspecification by considering non-genetic residual effects influenced by the environment. In response to the concerns observed, we propose novel GxE PRS models for both quantitative and binary traits, for which we have developed a user-friendly R package called “GxEprs”. To demonstrate the effectiveness of the proposed model, we applied it to analyze a diverse set of quantitative and binary traits using data from the UK Biobank. The results of our analyses showcase that the proposed GxE PRS model is robust against spurious GxE signals and addresses key limitations observed in existing models. This advancement contributes to improving the accuracy and reliability of GxE PRS modeling, providing researchers with a valuable tool to explore gene-environment interactions in the context of disease risk assessment.

## Methods

This section describes the existing and proposed statistical models for GxE PRS. Subsequently, it illustrates details on genotype data used for this study, along with the quality control procedures undertaken. Finally, it provides details on variables including phenotypes and covariates used in the real data analysis.

### Statistical Models

A conventional GWAS model in the discovery dataset can be written as,

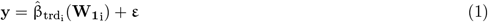

where **y** is the n_1_x1 vector of phenotypes that have been adjusted for all available fixed effects in the discovery sample,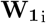 is the n_1_x1 vector of genotypes at the i^th^ SNP, 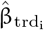 is the estimated additive effect of the i^th^ SNP, and **ε** denotes the n_1_x1 vector of residual terms, and n_1_ is the number of samples in the discovery dataset.

When considering GxE interactions, a GEWIS model can be written as,

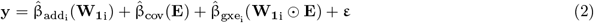

where **y** and 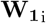 are as defined above, **E** is the n_1_x1 vector of the covariate that can modulate the genetic effects, 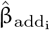, is the additive effect of the i^th^ SNP, 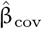 is the estimated effect of the covariate, 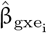 is the non-additive effect of GxE at the i^th^ SNP, and **ε** denotes the n_1_x1 vector of residual terms. Overall, this equation provides a means of examining the complex interplay between genetic and environmental factors in relation to the outcome variable.

In case there is a non-linear relationship between y and E because of various reasons such as underlying biological mechanisms or collider bias due to adjustment, we can use an alternative GWEIS model that includes quadratic effects of E. The model can be written as,

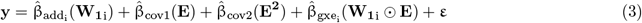

where **y**, 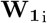, **E**, 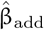 and 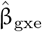 are as defined above, 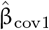 is the estimated effect of the covariate, 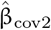 is the estimated effect of the squared covariate, and **ε** denotes the n_1_x1 vector of residual terms.

A conventional PRS which has been widely used to predict disease risk, calculated as the weighted sum of risk alleles identified in a GWAS. Based on GWAS summary statistics obtained from the discovery samples (eq.1), a set of PRSs can be constructed using the following model:

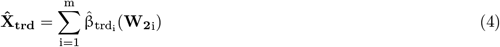

where 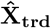 is the n_2_x1 vector of estimated PRSs, **W**_**2**i_ is the n_2_x1 vector of genotypes at the i^th^ SNP, m is the number of SNPs and n_2_ is the number of individuals in the target dataset.

Similarly using GWEIS summary statistics from eq. (2), two sets of PRSs in a target dataset can be estimated as follows:

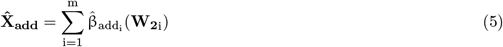

and

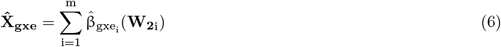

where 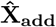 and 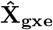 are the n_1_x1 vectors of estimated PRSs for additive and GxE interaction effects.

Using these three sets of PRSs estimated from eq. (4), (5) and (6), we will test five GxE models (referred to as Models 1 - 5) to test in the target dataset.

The first model, commonly used in the conventional GxE analysis, is written as,

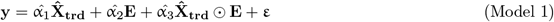

where **y** is the n_2_x1 vector of phenotypes adjusted for fixed effects in the target sample, **E** is the n_2_x1 vector of covariate, 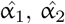 and 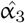 are the estimated regression coefficients of the PRS, covariate and interaction terms respectively, and **ε** denotes the n_2_x1 vector of residual terms of each target model.

The second model, slightly modified version of the conventional GxE model, is represented as follows:

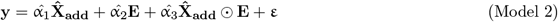

where **y, E** and **ε** are defined as above, 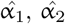 and 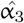 are the estimated regression coefficients of the PRS, covariate and interaction terms, respectively, modelled with 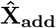 instead of 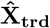.

The third model is similar to the second model, but uses 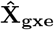 to model the interaction term, as follows:

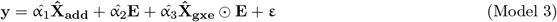

where variables are defined as above, and 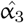 is the estimated regression coefficients of the interaction terms based on 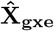 not 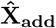. It is worth noting that this model has been previously used ^14,15^.

We propose two new extensions to the existing GxE PRS models, which are denoted as Models 4 and 5. Model 4 can be expressed as:

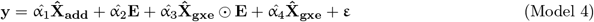

where variables are defined as in Model 3, and 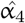 is the estimated regression coefficients for the PRS of GxE interaction 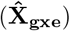. This model is more intuitive than Model 3 because the main effects of 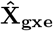 can be partitioned as spurious GxE interaction if not modelled.

To address the potential issue of spurious GxE signals that may arise from non-linear relationships between the outcome variable y and covariate E, we propose the inclusion of a quadratic polynomial component for E. This can be particularly relevant in cases of binary outcomes and sources of bias such as model misspecification. The proposed model, denoted as Model 5, is expressed as:

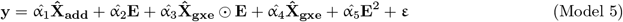

where variables are defined as in Model 4, and 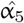 is the estimated regression coefficients for quadratic effects of E.

### Permutation

To address the potential issue of spurious GxE signals that may arise from unknown relationships between the quantitative outcome variable (**y**) and covariate (**E**), we implemented a permutation procedure on the term 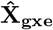 in the interaction component in Model 4 and denoted it as Model 4*. The purpose of this permutation was to maintain the correlation structure between the outcome variable and other model components while specifically permuting the interaction component. Similarly, in Model 5 for binary outcome, we performed the same permutation on the 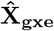 term in the interaction component and denoted it as Model 5*. It’s important to note that the number of permutations performed in Models 4* or 5* was determined based on the p value obtained in Models 4 or 5 to ensure an adequate number of permutations (with a minimum of 1000). By selectively permuting the interaction term while preserving the correlation structure, we aimed to assess the robustness of the GxE effects in our models and mitigate the risk of spurious signals. This approach allowed us to obtain more reliable results and evaluate the significance of the observed GxE interactions.

### Genotype Data and Quality Control Process in the UK Biobank

Genotype data were extracted from UK Biobank which is a population-based cohort of over 500,000 individuals from England, Scotland and Wales recruited between 2006 and 2010^19^. Genotyping was done using the UK Biobank Axiom Array with imputation using the Haplotype Reference Consortium and UK10K + 1000 Genomes reference panels ^20^. To minimize genetic heterogeneity, we only used the largest ancestry group; the White British population (408,183 individuals). Quality control (QC) procedures have been carried out for initial genotype data at the SNP level (INFO *<* 0.6, multi-character allele codes, minor allele frequency (MAF) *<* 0.01, the Hardy Weinberg Equilibrium (HWE) p *<* 10^−7^, SNP call rate *<* 0.95 and duplicate ID variants were excluded) and individual level (Non-White-British ancestry, missingness *>* 0.05, submitted and inferred sex mismatched, poor genotype quality or a sex chromosome aneuploidy ^21^ and closely related individuals where relationship *>* 0.05 based on Genetic Relationship Matrix (GRM) were excluded). A total of 7,701,772 SNPs and 288,792 individuals remained for further analyses. From these, HapMap3 SNPs were used in this study as they were considered robust and reliable for estimating heritability and genetic correlation ^22,23^, hence genetic prediction. After these quality controls, as genotype data, 1,217,311 (HapMap3) SNPs and 288,792 individuals remained for the analyses.

### Simulated Phenotype and Covariate Data

We used R ^24^ and MTG2^25^ software packages to simulate phenotypes of a complex trait, on a dataset consisting of 10,000 individuals and 10,000 SNPs, which were selected randomly from the 288,792 individuals and 1,217,311 SNPs. We generated three sets of SNP effects that were weighted to the real genotypes of the data, to simulate the genetic effects of the main phenotype (**G**_**0**_), the genetic effects of the environmental variable (**G**_**E**_) and the genetic effects interacted with the environmental variable (**G**_**1**_), using R library MASS ^26^.

To simulate phenotypes of a complex trait and evaluate the performance of the proposed GxE PRS models, we employed four different simulation models. The first is the null simulation model, where the phenotype is purely determined by the genetic effects of the main phenotype and random error, expressed as:

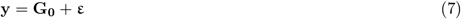

The second is the GxE simulation model, which includes the interaction effect between genetic effects and the environmental variable E, expressed as,

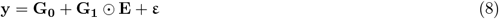

The third is the RxE simulation model, which considers the interaction effect between unknown random effects and the environmental variable **E** with absence of GxE, expressed as,

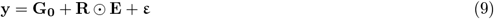

The fourth is the RxE simulation model, which considers the interaction effect between unknown random effects and the environmental variable **E** with presence of GxE, expressed as,

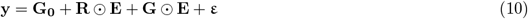

The fifth is the covariate (**E**) simulation model, which simulates the environmental variable **E** as a combination of genetic effects and random error, expressed as,

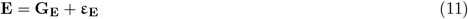

where **y** is the vector of simulated phenotypes, **G**_**0**_ is the vector of main genetic effects of the phenotype, **G**_**1**_ is the vector of genetic effects of the phenotype, **R** is the vector of random variables generated from either *N* (0, 0.25) or *N* (0, 0.5), **E** is the vector of environmental variable, **G**_**E**_ is the vector of genetic effects of **E, ε** is the vector of residuals of the main phenotypes and **ε**_**E**_ is the vector of residuals of **E**. Here, the genetic variances of the main phenotypes (Var(**G**_**0**_)) and the environmental variance (Var(**G**_**E**_)) were fixed to 0.4 and 0.5 respectively. The variance of interaction component (Var(**G**_**1**_)) was set to 0, 0.01 and 0.05 and covariance between **G**_**0**_ and **G**_**E**_ was varied among 0, 0.2, 0.4, 0.6 and 0.8. The residual structure was defined such that, residual variance of phenotype (**ε**) and residual variance of environmental variable (**ε**_**E**_) were both fixed to 0.5 and the residual covariance between the phenotype **y** and the environmental variable **E** was varied among 0, 0.2, 0.4, 0.6 and 0.8. Finally, separate data files for the main phenotypes and environmental variable (covariate) were stored for the downstream analysis.

To generate binary outcome variables, we followed the same process as described above for quantitative traits except that case-control status was determined based on the liability threshold model, given a predefined population prevalence rate. Specifically, we first simulated quantitative continuous phenotypes using the method outlined earlier. Next, using a population prevalence of 1%, 10%, or 50%, we determined the threshold for liability, using standard normal distribution theory, to classify individuals as cases or controls based on their quantitative continuous phenotypes. Covariates were generated using the same method as for the simulation of quantitative continuous phenotypes, using eq. (11).

In addition, we consider the possibility of RxE interaction effects generated by unmeasured or unobserved factors, as shown in eq. (9). These interactions can be examined by GxE PRS Models 1 – 5, but they can also result in spurious GxE due to model misspecification problem. Model misspecification occurs when the statistical model used to describe the data does not accurately represent the underlying data-generating process. This can happen due to a variety of reasons, such as incorrect functional form, omitted variables, or unmeasured or unknown factors that contribute to the variation in the response variable. It is a common issue in statistical modelling that can lead to biased estimates and incorrect inferences. Therefore, we explicitly assessed GxE PRS Models 1 – 5 for this potential model misspecification issue.

### Real Data

In real data analyses, our primary objective was to demonstrate how these models can be applied in real-world scenarios. We aimed to showcase how these GxEPRS models can be used to identify signals for existence of GxE.

#### Sample Sizes

After performing QC (see section “Genotype Data and Quality Control Process” for details), 288,792 individuals remained for the analyses. We randomly split the total number of individuals into 2 datasets in the ratio of 8:2, namely discovery and target datasets. The discovery dataset was used to perform GWAS/GWEIS while the target dataset was used to predict risk scores.

#### Real Phenotypic Data

Real phenotypic data were extracted from the UK Biobank. The phenotypes include anthropometrical traits such as body mass index (BMI), waist-to-hip ratio (WHR), and clinical traits such as low-density lipoprotein (LDL) cholesterol, which are quantitative traits. In addition, we considered binary traits such as diabetes (DIAB), hypertension (HYP) and coronary artery disease (CAD).

#### Covariates

To account for non-genetic environmental effects, we included fixed effects such as sex, age at recruitment, Townsend deprivation index (TDI), education in years ^27^, and the first 10 genetic principal components ^28^ in our analysis. These variables were incorporated to adjust for main phenotypes and control for confounding factors. In the discovery dataset, the phenotype (outcome) was adjusted during the GWAS/GWEIS stage. In the target dataset, these confounders were included in the respective target models to capture their effects on the outcome.

In the GxE analysis, we considered several environmental variables. For quantitative outcomes (BMI, WHR, or LDL), we examined three covariates: frequency of alcohol consumption (ALC), healthy diet (HD), and physical activity (PA) as the **E** variable in GxE. For binary outcomes (CAD, DIAB, or HYP), we examined four covariates: BMI, high-density lipoprotein (HDL), haemoglobin (HGB), and WHR as the **E** variable in GxE. Each outcome was analyzed separately with its corresponding covariates. It is important to note that these covariates were standardized independently in the discovery and target datasets before being used in the downstream analysis.

For a comprehensive overview of the variables used in our study, including detailed information on the adjusted phenotypes and environmental variables, please see Tables S1, S2 and S3.

## Results

### Quantitative Traits

Figure 1 presents the summarized results of the simulation study for quantitative traits. The study included varying levels of genetic and residual correlations, with Cor(**G**_**0**_, **G**_**E**_) values set at 0, 0.4, and 0.8, and Cor(**ε, ε**_**E**_) values set at 0, 0.4, and 0.8. In the discovery dataset, SNP effects were estimated using eq. (1) (GWAS) and eq. (2) (GWEIS). In the absence of GxE interaction in the simulations (Var(GxE)=0), Models 1 – 4 were controlled for a type 1 error rate of 5% (Figure 1 a). However, in the presence of GxE interaction in the simulation (Var(GxE)=0.01 or 0.05), the statistical power of Models 3 and 4 outperformed both Models 1 and 2. Note that Model 1 – 3 are existing models while Model 4 is newly introduced in this study. When comparing Models 3 and 4, there is no remarkable difference in their performance. Figure S1 provides more results for additional data points in regard to genetic and residual correlations.

**Figure 1:**
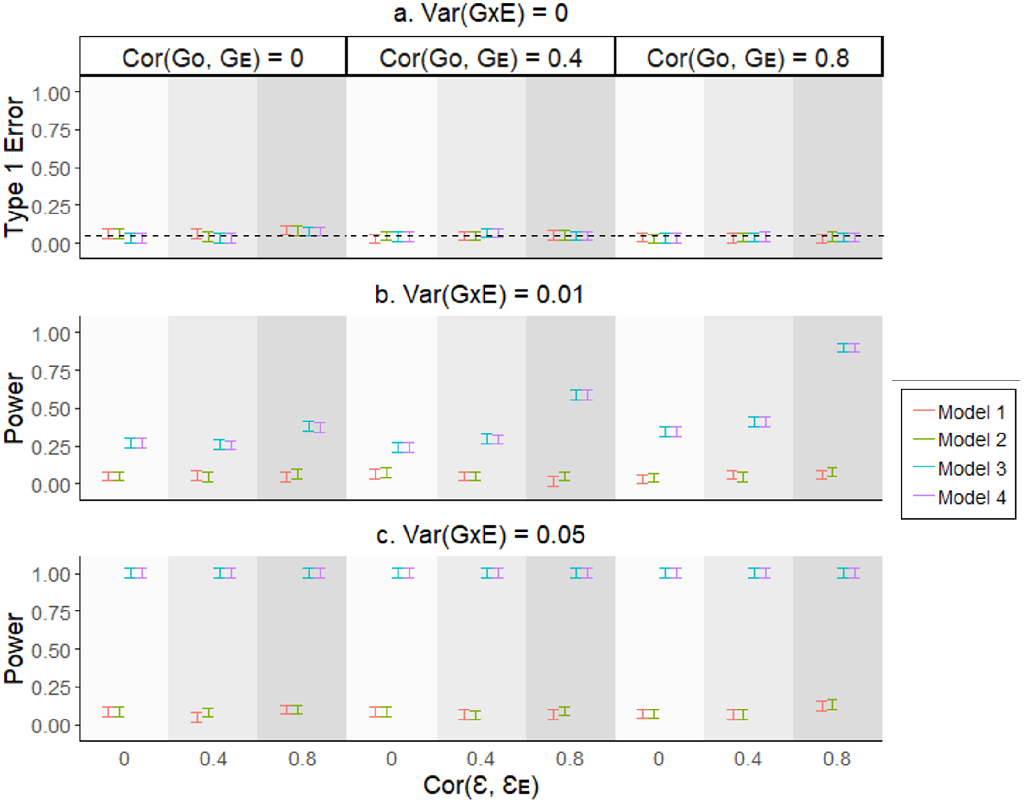
The type 1 error rate and statistical power of various GxE PRS models when using quantitative traits. We used simulation models (eq. 7, 8, and 11) to generate phenotypes and covariates. Different genetic and residual correlations (Cor(**G**_**0**_, **G**_**E**_) = 0, 0.4 and 0.8), and Cor(**ε, ε**_**E**_) = 0, 0.4, and 0.8) were considered in various scenarios. We applied Models 1 - 4 to estimate the GxE component, with SNP effects estimated from GWAS (eq. 1) or GWEIS (eq. 2). The error bars show the 95% confidence intervals for type 1 error rate and statistical power (vertical axes), based on averaging results from 200 simulated replicates.

### Binary Traits

Figure 2 presents the simulation study results for binary traits with a population prevalence of 10%. Similar to the simulation for quantitative traits (Figure 1), we varied levels of genetic and residual correlations, and estimated SNP effects using eq. (1) (GWAS) and eq. (2) (GWEIS). When using binary traits, the existing methods (Models 1 - 3), showed inflated type 1 error rates, particularly at higher genetic and/or residual correlations (refer to Figure 2 a). However, the proposed method (Model 4), maintained a well-controlled type 1 error rate at 5% in all cases. Regarding statistical power, Model 4 outperformed Models 1 and 2, exhibiting higher power in the presence of large GxE effects (see Figure 2 c, d and e). It is important to note that although Model 3 showed higher power, this may be due to type 1 error rate inflation (as evidenced in Figure 2 a). When GxE was low (i.e., 0.01), there appeared to be no power to detect GxE using Model 4 (see Figure 2 b). This may be due to overall power decrease when using binary responses (as shown in Figure 1 vs. 2). For additional data points in regards to genetic and residual correlations, please refer to

**Figure 2:**
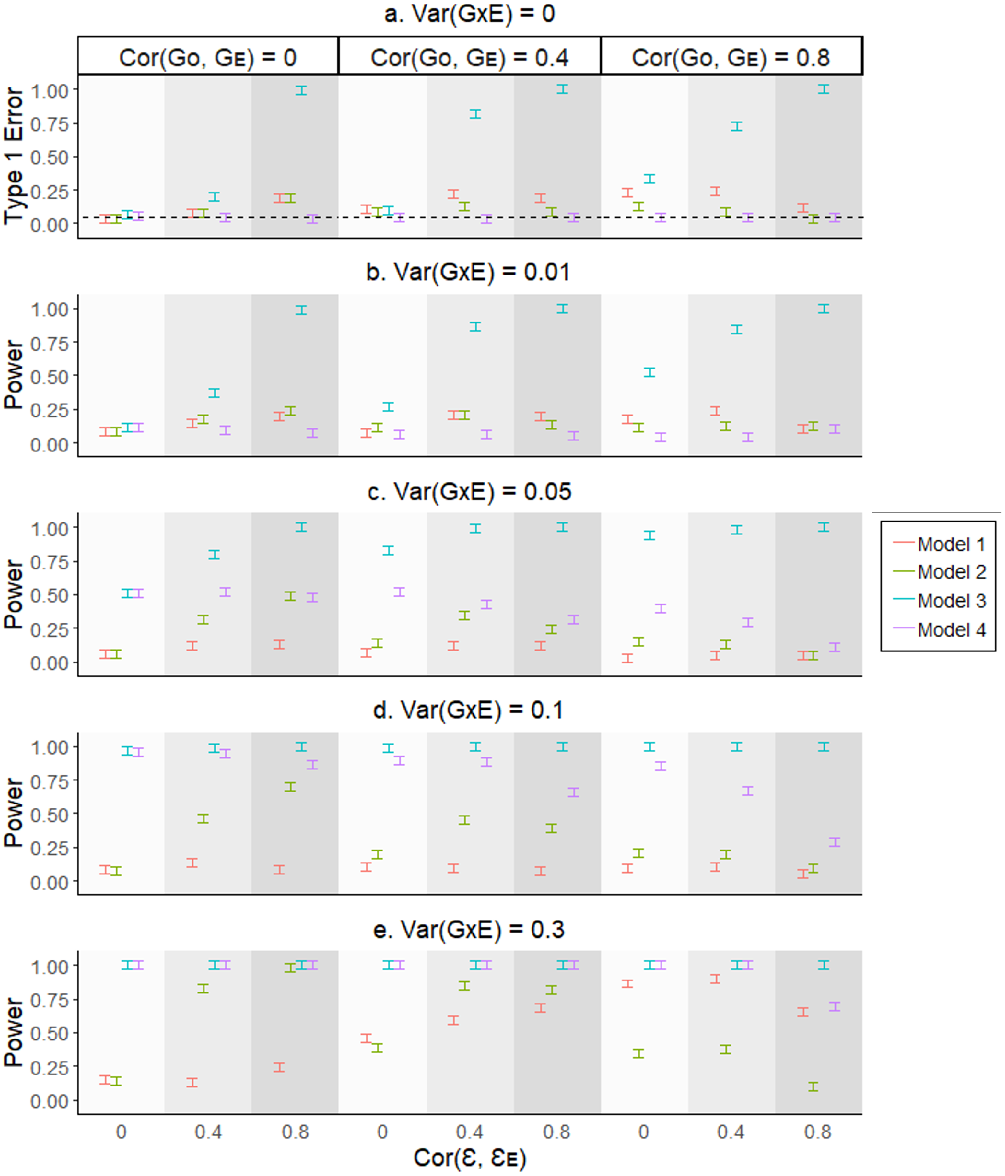
The type 1 error rate and power comparison of existing and proposed methods for binary traits with 10% population prevalence. To assess the performance of our proposed method (Model 4) and existing methods (Models 1-3) on binary traits, we conducted simulation studies using various scenarios. We generated quantitative phenotypes and covariates using simulation models (eq. 7, 8, and 11) and converted them to binary phenotypes using a liability threshold of 10% population prevalence. We considered different levels of genetic and residual correlations (Cor(**G**_**0**_, **G**_**E**_) = 0, 0.4 and 0.8), and Cor(**ε, ε**_**E**_) = 0, 0.4, and 0.8) and estimated SNP effects using GWAS (eq. 1) or GWEIS (eq. 2). We applied Models 1-4 to estimate the GxE component. The error bars in the figure represent the 95% confidence intervals for type 1 error rate and statistical power (vertical axes), and are based on averaging results from 200 simulated replicates.

Figure S2. Moreover, when we tested different population prevalence values (1% and 50%), we consistently observed that Model 4 controlled the type 1 error rate and exhibited reasonable statistical power, unless var(GxE) was negligible (see Figures S3 and S4). However, Models 1-3 either showed inflated type 1 error rates or lacked power.

#### Assessing the Impact of Model Misspecification on Quantitative Traits in GxE PRS Models

To evaluate the potential issue of model misspecification in GxE PRS models, we assessed Models 1-5 using a simulation analysis for quantitative traits. The true simulation models were defined by eq. (9), (10) and (11), and a range of values were selected for genetic and residual correlations (Cor(**G**_**0**_, **G**_**E**_) = 0, 0.4, and 0.8, and Cor(**ε, ε**_**E**_) = 0, 0.4, and 0.8), and RxE interaction effects generated by unmeasured or unobserved factors (Var(RxE) = 0.25 or 0.5). Figure 3 a illustrates that Models 1 – 4 generated an inflated type I error rate due to model misspecification. However, to remedy this inflation issue, a permutation technique was found to be useful (i.e. Model 4*) as shown in Figure 3 a.

**Figure 3:**
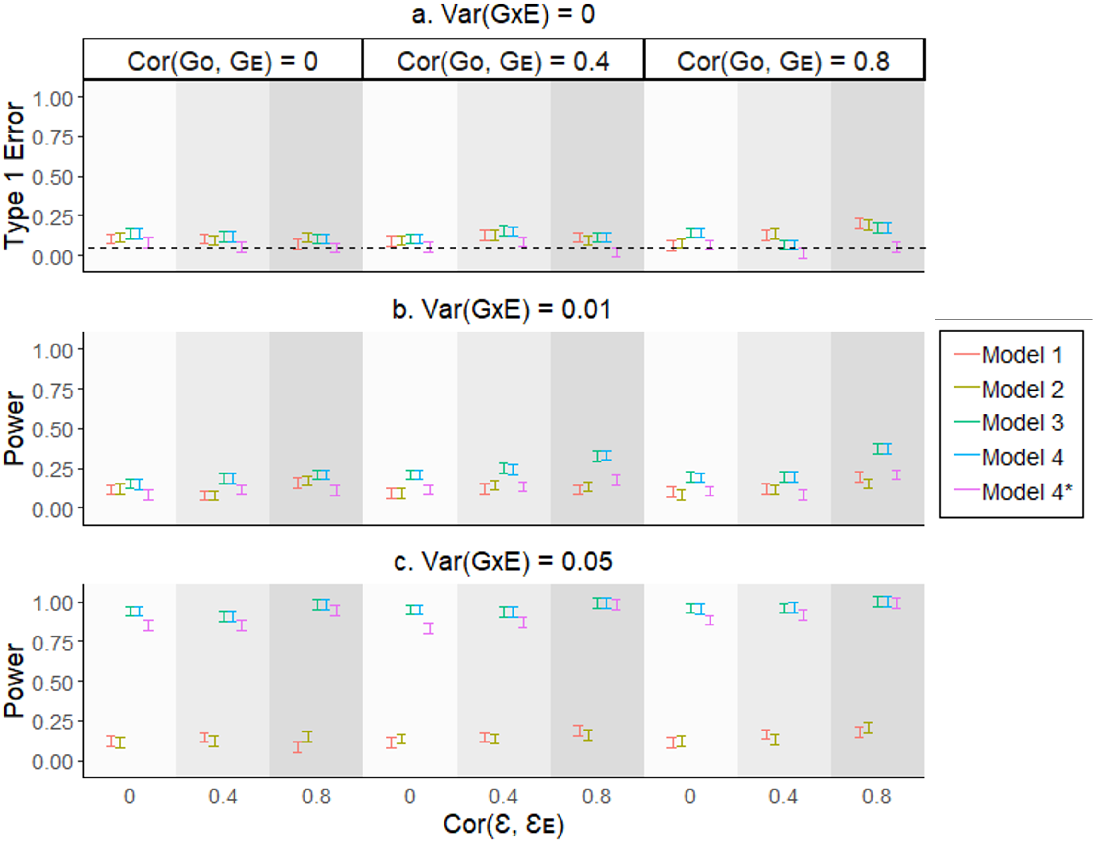
The type 1 error rate and statistical power of various GxE PRS models when using quantitative traits with Var(RxE) = 0.25. We used simulation models (eq. 9, 10, and 11) to generate phenotypes and covariates. Different genetic and residual correlations (Cor(**G**_**0**_, **G**_**E**_) = 0, 0.4 and 0.8), and Cor(**ε, ε**_**E**_) = 0, 0.4, and 0.8) were considered in various scenarios. Var(RxE) was set to 0.25. In the absence of GxE (eq. 9), we simulated the quantitative trait by adding the RxE component with the residual term. In the presence of GxE (eq. 10), we added the GxE component in addition to RxE and residual terms to simulate the quantitative trait. We applied Models 1 - 4 to estimate the GxE component, with SNP effects estimated from GWAS (eq. 1) or GWEIS (eq. 2). Model 4* is the permuted version of Model 4, obtained by permuting the 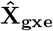 term of 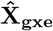 ⊙**E** component 1000 times. The error bars show the 95% confidence intervals for type 1 error rate and statistical power (vertical axes), based on averaging results from 200 simulated replicates.

In the presence of GxE, Models 1 and 2 demonstrated a lack of power even with Var(GxE) set to 0.05 as illustrated in Figures 3 b and c. On the other hand, Models 3, 4, and 4* demonstrated reasonable power unless Var(GxE) was negligible, as depicted in Figure 3 c. Figures S5 and S6 provide more results for additional data points in regards to genetic and residual correlations when using Var(RxE) = 0.25 and Var(RxE) = 0.5 respectively.

#### Assessing the Impact of Model Misspecification on Binary Traits in GxE PRS Models

To assess the impact of model misspecification in GxE PRS models for binary traits, we conducted a simulation analysis using Models 1-5 with a population prevalence of 10% (Figure 4). The true simulation models were defined by eq. (9), (10) and (11), and binary responses were generated using a liability threshold model. However, in the absence of an interaction term (Var(GxE) = 0), none of the models, including the permutation model (Model 4*), controlled the type 1 error rate, regardless of whether Var(RxE) was 0.25 or 0.5 (Figure 4 and Figure S7).

**Figure 4:**
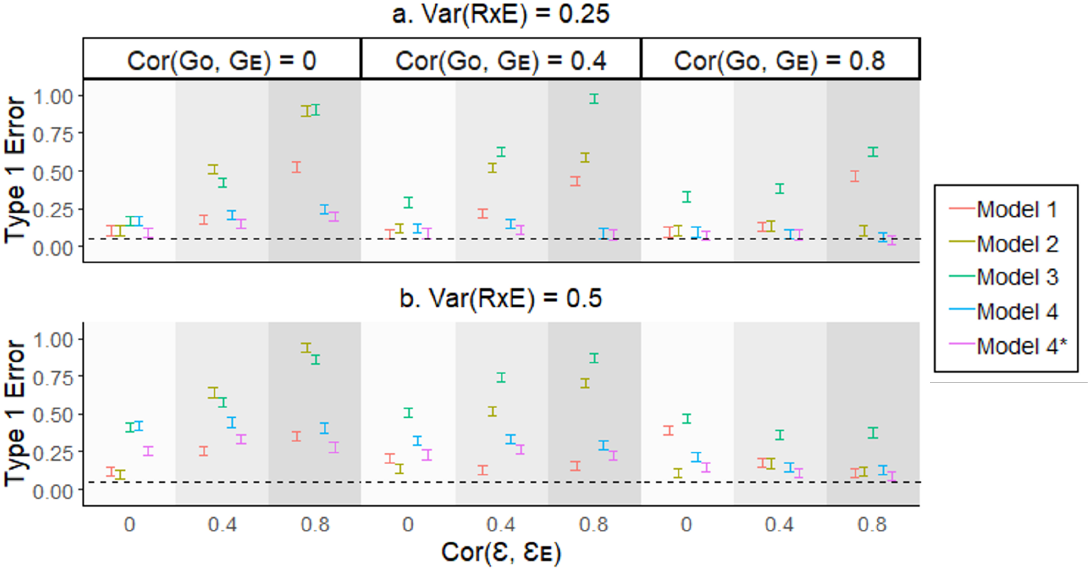
The type 1 error rate of various GxE PRS models when using binary traits with Var(RxE) = 0.25 or 0.5. We used simulation models (eq. 9, 10, and 11) to generate phenotypes and covariates. We used liability threshold of 10% population prevalence to simulate binary phenotypic outcomes. Different genetic and residual correlations (Cor(**G**_**0**_, **G**_**E**_) = 0, 0.4 and 0.8), and Cor(**ε, ε**_**E**_) = 0, 0.4, and 0.8) were considered in various scenarios. In the absence of GxE (eq. 9), we simulated the quantitative trait by adding the RxE component with the residual term, and then converted to binary scale using liability threshold of 10% population prevalence. We applied Models 1 - 4 to estimate the GxE component, with SNP effects estimated from GWAS (eq. 1) or GWEIS (eq. 2). Model 4* is the permuted version of Model 4, obtained by permuting the 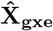 term of 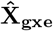 ⊙**E** component 1000 times. The error bars show the 95% confidence intervals for type 1 error rate and statistical power (vertical axes), based on averaging results from 200 simulated replicates.

To address the issue of model misspecification, we further adjusted the proposed model by incorporating a second-order covariate term (**E**^2^) in both discovery and target models (Model 5). When we applied Model 5 with the adjusted GxE PRS method, we observed effective control of the type 1 error rate at 5%, regardless of whether Var(RxE) was 0.25 or 0.5 (Figure 5 a, Figures S8 a, Figures S9 a). Model 5 also provided reasonable power when increasing the variance of GxE interaction (Figure 5 b, c, and d, Figure S8 b, c, and d, Figure S9 b, c, and d).

**Figure 5:**
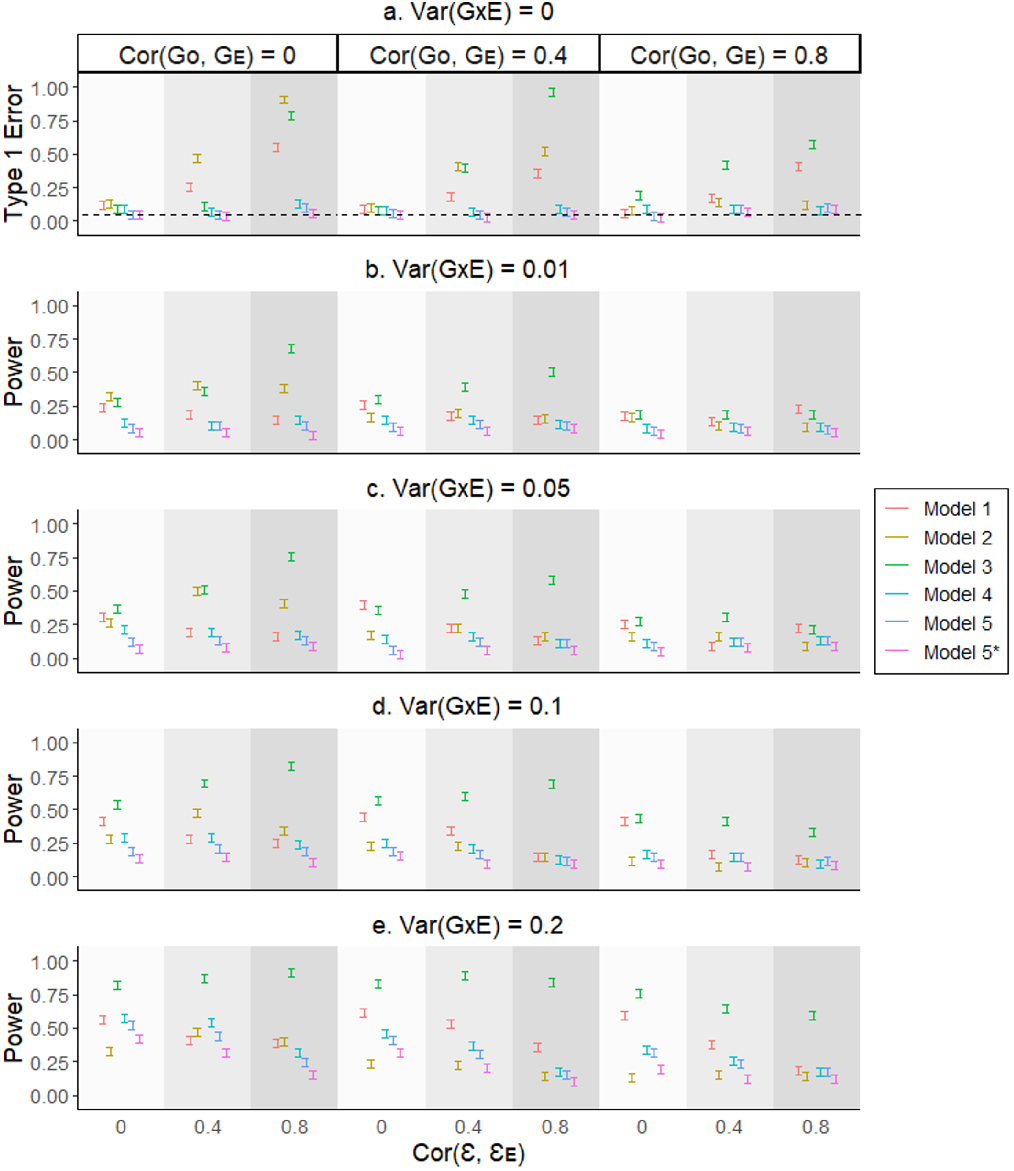
The type 1 error rate and power of various GxE PRS models when using binary traits with Var(RxE) = 0.25 and 10% population prevalence. To investigate the impact of model misspecification, we generated phenotypes and covariates with RxE effects using simulation models (eq. 9 and 10). The genetic and residual variances were set to fixed values of 0.4 and 0.5, respectively, for the main phenotypes. We used liability threshold of 10% population prevalence to simulate binary phenotypic outcomes. Different genetic and residual correlations (Cor(**G**_**0**_, **G**_**E**_) = 0, 0.4 and 0.8), and Cor(**ε, ε**_**E**_) = 0, 0.4, and 0.8) were considered in various scenarios. Var(RxE) was set to 0.25. In the absence of GxE (eq. 9), we simulated the quantitative trait by adding the RxE component with the residual term, and then converted to binary scale using liability threshold of 10% population prevalence. In the presence of GxE (eq. 10), we added the GxE component in addition to RxE and residual terms to simulate the quantitative trait and converted to binary scale in a similar manner. We applied Models 1 - 5, with SNP effects estimated from GWAS (eq. (1)) or GWEIS (eq. (3)). Model 5* is the permuted version of Model 5, obtained by permuting the 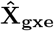 term of 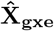 ⊙**E** component 1000 times. The error bars represent the 95% confidence intervals of the type 1 error rate and statistical power (vertical axes), based on averaging the results from 200 simulated replicates.

Furthermore, we investigated the influence of varying population prevalence, specifically at 1% or 50%, on the evaluation of the models. When the population prevalence was set to 1%, Models 5 and 5* effectively controlled their type 1 error rate (see Figures S10 a and S11 a). As the population prevalence increased to 50%, all the models maintained control over their type 1 error rate (see Figures S12 a and S13 a). However, when considering the presence of GxE interactions, Model 5 did not achieve sufficient statistical power at a population prevalence of 1%. It is worth noting that Model 1 appeared to have relatively higher power at lower levels of genetic and residual correlations, but this observation may be attributed to the inflated type 1 error rate of Model 1 (see Figures S10 and S11). In contrast, when the population prevalence was increased to 50, Models 3-5* demonstrated reasonable statistical power, especially when the variance of the GxE interaction was large (see Figures S12 and S13).

### Real Data Analysis

We analyzed BMI, LDL and WHR as quantitative traits with relevant modifiable covariates such as ALC, HD and PA across each GxE PRS model including Models 1 - 4 where Model 4 is the proposed model (see “Methods” section). We recorded the estimates and significance of the interaction term of interest (i.e. PRSxE component in the prediction model). We aimed to identify the significant GxE effects from the proposed models and to compare the PRSxE estimates and the corresponding significance values across all the target models, Models 1-4 and 4* (the permuted model of Model 4).

In Figure 6, it is evident that the genetic effects of BMI are modulated by ALC, as indicated by the significance of Models 4 and 4*. However, although Models 1 and 2 identified a significant GxE effect in the analyses of the outcome/covariate pair BMI/PA, the significance was not confirmed by the proposed models (Models 4 and 4*). Considering the limitations of Models 1 to 3 in controlling the type 1 error rate, as demonstrated in the simulation study, we cannot solely rely on the GxE effects identified by Models 1 and 2 (Figure 6). Regarding the analysis of WHR/ALC, a significant GxE effect was observed in Model 4*, which is the permutation of Model 4 and designed to address spurious signals. It is important to note that Models 4 and 4* yield similar p values, both of which are very close to the significance threshold (refer to Table S10). Therefore, based on the findings from Figure 6 and considering the limitations of Models 1 - 3 in controlling the type 1 error rate, we conclude that, except for the BMI/ALC and WHR/ALC outcome/covariate pairs, there is insufficient evidence to support the presence of GxE effects for the traits assessed.

**Figure 6:**
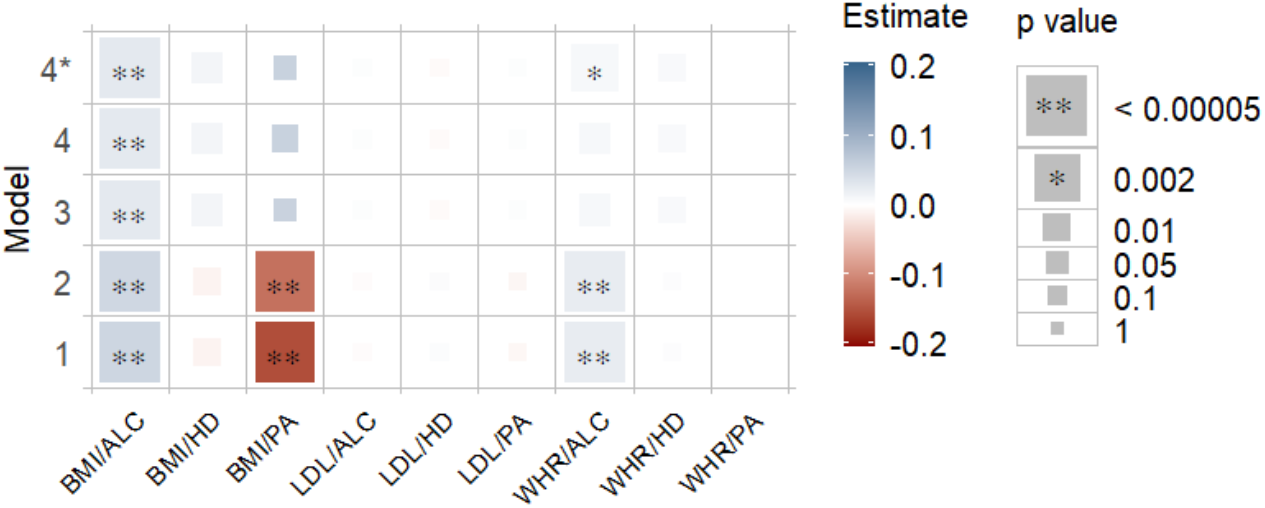
Estimates and Significance of PRSxE Components for Quantitative Phenotypes across Models The heatmap represents the estimated regression coefficient of the PRSxE term for each model. The color gradient ranges from dark red to dark blue, indicating the magnitude of the regression coefficient. The size of each square in the heatmap is proportional to the corresponding p value. Significance levels are indicated by asterisks, representing the significance after Bonferroni correction (significance level = 0.05/21 ∼ 0.002), considering a total of 21 analyses conducted. The number of permutations performed in Model 4* was determined based on the p value obtained in Model 4 to ensure an adequate number of permutations (with a minimum of 1000). The horizontal axis represents the outcome/covariate pairs, while the vertical axis represents the models used for estimating GxE.

We also analyzed CAD, DIAB and HYP as binary traits with relevant modifiable covariates such as BMI, HDL, HGB and WHR across each GxE PRS model including Models 1 - 5 where Model 5 is the proposed model (see “Methods” section). We recorded the estimates and significance of the interaction term of interest (i.e. PRSxE component in the prediction model). We aimed to identify the significant GxE effects from the proposed models and to distinguish the PRSxE estimate and the corresponding significance values amongst all the target models (Models 1-5 and 5*. Here Model 5* is the permuted model of Model 5). The results are as described below.

Figure 7 illustrates that the genetic effects of HYP are modulated by WHR, as supported by the significance of Models 5 and 5*. Model 3 identified a significant GxE effect in the analyses of CAD/HDL, DIAB/BMI, and DIAB/HDL, but these were not confirmed by Models 5 and 5*. Similarly, Models 1 or 2 identified significant GxE effects in the analyses of CAD/BMI, CAD/HGB, and DIAB/HGB, but these were not confirmed by Models 5 and 5*. Considering that Models 1 to 4 lack control over the type 1 error rate for binary diseases, as demonstrated in the simulation verification, we cannot solely rely on the GxE effects identified by Model 1, 2, or 3 (Figure 7). However, it is noteworthy that HYP/WHR exhibited significant GxE estimates when Models 5 and 5* were employed. Therefore, based on the findings from Figure 7, we conclude that there is no reliable evidence for GxE effects in the binary outcome/covariate pairs, except for HYP/WHR.

**Figure 7:**
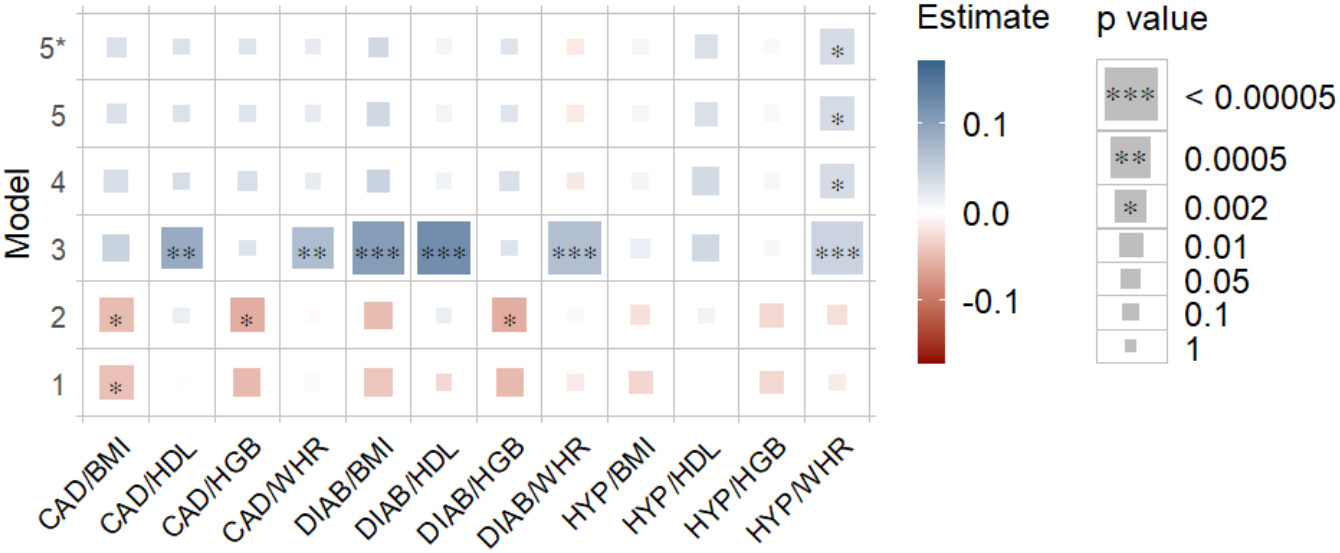
Estimates and Significance of PRSxE Components for Binary Phenotypes across Models The heatmap represents the estimated regression coefficient of the PRSxE term for each model. The color gradient ranges from dark red to dark blue, indicating the magnitude of the regression coefficient. The size of each square in the heatmap is proportional to the corresponding p value. Significance levels are indicated by asterisks, representing significance after Bonferroni correction (significance level = 0.05/21 ∼ 0.002), considering a total of 21 analyses conducted. The number of permutations performed in Model 5* was determined based on the p value obtained in Model 5 to ensure an adequate number of permutations (with a minimum of 1000). The horizontal axis represents the outcome/covariate pairs, while the vertical axis represents the models used for estimating GxE.

## Discussion

The present study aimed to evaluate the performance of various GxE PRS models for both quantitative and binary traits, develop an improved modelling approach that is more robust and assess the impact of genetic and residual correlations between the main phenotypes and covariates (i.e., **E**) as well as model misspecification ^16^. The results obtained from this study provide important insights into the effectiveness and limitations of GxE PRS models ^16^.

For quantitative traits, Models 3 and 4 demonstrated superior performance compared to Models 1 and 2 in terms of controlling type 1 error rates and statistical power to detect GxE interactions. Model 4, introduced in this study, performed similarly to Model 3, indicating its efficacy in capturing GxE interactions. However, in the presence of model misspecification, none of the GxE PRS models except the permutation of Model 4 worked effectively. This highlights the importance of carefully selecting GxE PRS models when detecting GxE interactions.

For binary traits, existing models (Models 1-3) exhibited inflated type 1 error rates, particularly at higher levels of genetic and/or residual correlations. However, the proposed Model 4 maintained a well-controlled type 1 error rate of 5% in all scenarios. Additionally, Model 4 displayed higher power compared to Models 1 and 2 in the presence of GxE effects. It is worth noting that Model 3 showed higher power, but this may be attributed to its inflated type 1 error rate. To address the issue of model misspecification, the adjusted Model 5, which incorporated a second-order covariate term (**E**^**2**^) in both the discovery and target models, effectively controlled the type 1 error rate regardless of the magnitude of the variance of the source for model misspecification (e.g., RxE interaction).

The analysis of real data further supported the efficacy of the proposed models. Given an outcome/covariate pair, a variation in significant results across models justifies this point by agreeing with simulation findings. That is, we were capable of correctly identifying, whether the existence of GxE effects was true or not, in the instances where existing models resulted in spurious signals. For quantitative traits, significant GxE effects were identified in the analysis of BMI/ALC ^29,30^ and WHR/ALC, confirmed by the permuted version of Model 4 (Model 4*). For binary traits, high significance was found in the PRSxE component in the HYP/WHR analysis ^30,31^. These findings highlight the importance of considering GxE interactions in prediction models and demonstrate the potential of GxE PRS models for uncovering gene-environment interplay in complex traits and diseases.

Not all GxE interaction found in this study align with previous studies ^29,32,30^, including inconsistencies with our own previous studies ^29,30^. There are several potential reasons for this discrepancy. Firstly, previous studies relied on traditional models based on the discovery dataset only, without incorporating GxE PRS models like the current study. By using GxE PRS models and incorporating both the discovery and target datasets, our study may yield more robust outcomes. Secondly, previous studies, which employed GxE PRS models, mostly used Model 1, 2 or 3. Consequently, the observed signals in those studies may have been influenced by factors such as lack of power, genetic and residual correlation, and potential model misspecification. In contrast, our study adopted a novel approach (Models 4 and 5) that addresses these factors, enhancing the accuracy and reliability of our results. Lastly, it is worth considering that the study-wise Bonferroni correction employed in this study may have led to over-correction, potentially diminishing the significance of certain signals. Therefore, a cautious interpretation of the results is necessary.

Overall, the results of this study emphasize the significance of GxE interactions in the context of PRS models and provide valuable insights into the performance and limitations of various GxE PRS models. The findings support the potential utility of incorporating GxE effects to improve the accuracy and power of genetic risk prediction models for both quantitative and binary traits. However, it is important to carefully consider model misspecification issues ^16^ and potential confounding factors ^14,15^ when applying GxE PRS models in real-world settings.

One limitation of our study is that we cannot guarantee the directionality of the GxE model. For example, in the BMI/ALC analysis, we did not explicitly test the reverse direction, where the genetic effects of ALC are modulated by BMI status, due to limitations in our approach. Further research is needed to explore and determine the causal direction in the context of GxE interactions ^30,15^. Additionally, there are other factors that may influence the performance of GxE PRS models, such as variations in population prevalence and the interplay between multiple environmental factors. Moreover, investigating the robustness and generalizability of these models across diverse populations and traits would be valuable in advancing our understanding of GxE interactions in complex phenotypes.

In conclusion, this study provides valuable insights into the performance and applicability of GxE PRS models for both quantitative and binary traits. The findings support the importance of considering GxE interactions in genetic risk prediction and highlight the potential of GxE PRS models for uncovering complex gene-environment interplay. The proposed models are useful to study GxE interactions and their future applications have implications for personalized medicine and risk assessment, as they provide a framework for incorporating environmental factors into genetic risk prediction models.

## Acknowledgements

This research is supported by the Australian Research Council (DP190100766). D.J. acknowledges funding from the Australian Government Research Training Program (RTP) scholarship. We obtained real data from UK Biobank and the authors would like to acknowledge for providing access for the data sources. Our reference number approved by UK Biobank is 14575. The UK Biobank is funded by the UK Department of Health, the Medical Research Council, the Scottish Executive, and the Welcome Trust medical research charity. The analyses were performed using computational resources provided by the Australian Government through Gadi under the National Computational Merit Allocation Scheme (NCMAS) and under University of South Australia (UniSA), and HPCs (Statgen and Statgen 2 servers) managed by UniSA IT. We thank University of South Australia IT team for their support in accessing the servers. Finally we would like to thank the Statistical Genetics Group at the Australian Center for Precision Health for their support in providing quality controlled genotypic and phenotypic data.

## Author Contributions

S.H.L conceived the idea, derived the proposed GxE PRS models, and supervised the study. D.J. contributed in performing simulations, data extraction, data analysis and visualizations. D.J and S.H.L wrote the first draft of manuscript. D.J. and M.M.M. worked in R package preparation. S.H.L. initially created computer codes for the simulations and analyses, and reviewed simulations, data extraction, data analysis and visualizations throughout the study. M.M.M., K.B., E.H. and B.B. reviewed the manuscript and provided critical feedback with suggestions. All the authors discussed the results and contributed to finalizing the manuscript.

## Software Availability

The GxEprs source code, manual and example files can be downloaded from GitHub https://github.com/DoviniJ/GxEprs

