## Supplementary file for "GxE PRS: Genotype-environment interaction in polygenic risk score models for quantitative and binary traits"

### Supporting Information

#### Quantitative Traits

Figure S1 presents the results of the simulation study for quantitative traits. The study involved varying levels of genetic and residual correlations, with Cor($\mathbf{G_0}$, $\mathbf{G_E}$) values set at 0, 0.2, 0.4, 0.6 and 0.8, and Cor($\boldsymbol{\varepsilon}$, $\boldsymbol{\varepsilon_E}$values set at 0, 0.2, 0.4, 0.6 and 0.8. In the discovery dataset, SNP effects were estimated using eq. (1) (GWAS) and eq. (2) (GWEIS). All four models were controlled for a type 1 error rate of 5%. However, in terms of statistical power, Models 3 and 4 outperformed both Model 1 and Model 2, particularly when considering the impact of gene-environment interactions (GxE) with Var(GxE) set at 0.01 and 0.05. These findings suggest that Models 3 and 4 are more effective in capturing the complexities of the genetic and environmental factors underlying quantitative traits.

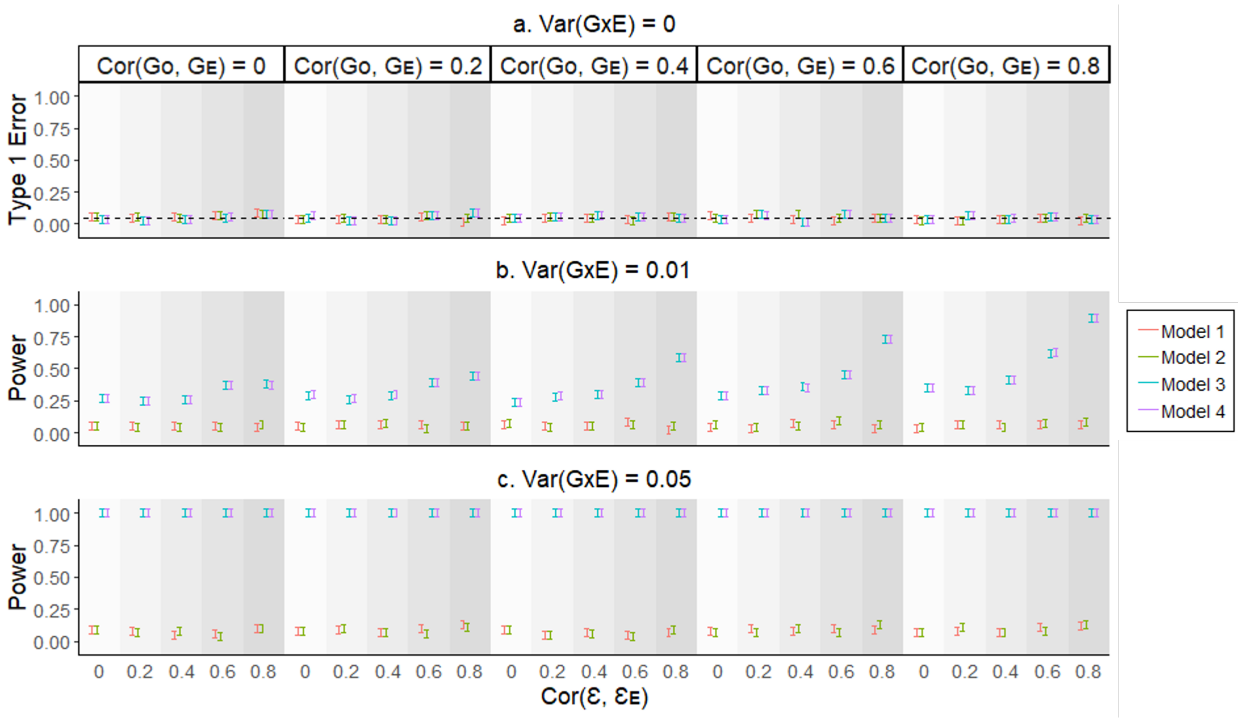

Figure S1: The type 1 error rate and statistical power of various GxE PRS models when using quantitative traits.

We used simulation models (eq. 7, 8, and 11) to generate phenotypes and covariates. Different genetic and residual correlations (Cor($\mathbf{G_0}$, $\mathbf{G_E}$) = 0, 0.2, 0.4, 0.6 and 0.8), and Cor($\boldsymbol{\varepsilon}$, $\boldsymbol{\varepsilon_E}$) = 0, 0.2, 0.4, 0.6 and 0.8) were considered in various scenarios. We applied Models 1 - 4 to estimate the GxE component, with SNP effects estimated from GWAS (eq. 1) or GWEIS (eq. 2). The error bars show the 95% confidence intervals for type 1 error rate and statistical power (vertical axes), based on averaging results from 200 simulated replicates.

#### Binary Traits

Figure S2 presents the results of the simulation study for binary traits with a population prevalence of 10%. The study involved varying levels of genetic and residual correlations, with Cor($\mathbf{G_0}$, $\mathbf{G_E}$) values set at 0, 0.2, 0.4, 0.6 and 0.8, and Cor($\boldsymbol{\varepsilon}$, $\boldsymbol{\varepsilon_E}$values set at 0, 0.2, 0.4, 0.6 and 0.8. In the discovery dataset, SNP effects were estimated using eq. (1) (GWAS) and eq. (2) (GWEIS). Model 1 has an inflated type 1 error rate especially at higher genetic correlation values (see Figure S2 a). When there is no genetic and residual correlation or no genetic and very low residual correlation (i.e. 0.2), type 1 error rate of Model 1 is controlled. Type 1 error rate of Model 2 possesses an unclear pattern, unlike Model 1. However, type 1 error rate of Model 3 is always controlled (at 5%). With respect to the statistical power, it is clear that Model 3 outperforms Models 1 and 2, where the power is higher in the presence of high GxE (see Figure S2 c and d). Power in Model 4 is relatively lower than the Models 1 and 2 in the presence of low GxE (see Figure S2 b), but the difference between the three models are negligible.

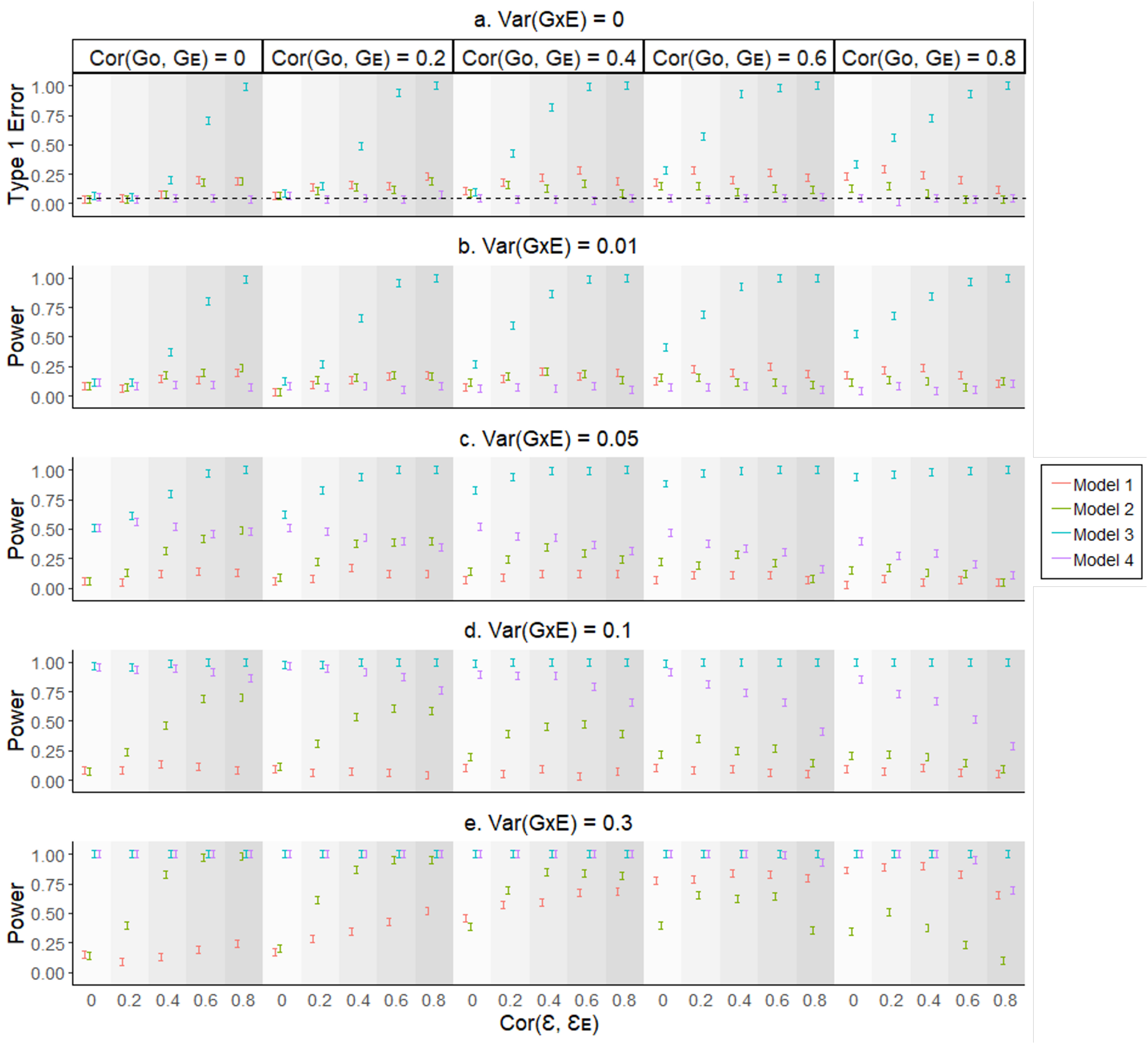

Figure S2: The type 1 error rate and power comparison of existing and proposed methods for binary traits with 10% population prevalence.

To assess the performance of our proposed method (Model 4) and existing methods (Models 1-3) on binary traits, we conducted simulation studies using various scenarios. We generated quantitative phenotypes and covariates using simulation models (eq. 7, 8, and 11) and converted them to binary phenotypes using a liability threshold of 10% population prevalence. We considered different levels of genetic and residual correlations (Cor($\mathbf{G_0}$, $\mathbf{G_E}$) = 0, 0.2, 0.4, 0.6 and 0.8), and Cor($\boldsymbol{\varepsilon}$, $\boldsymbol{\varepsilon}_\mathbf{E}$) = 0, 0.2, 0.4, 0.6 and 0.8) and estimated SNP effects using GWAS (eq. 1) or GWEIS (eq. 2). We applied Models 1-4 to estimate the GxE component. The error bars in the figure represent the 95% confidence intervals for type 1 error rate and statistical power (vertical axes), and are based on averaging results from 200 simulated replicates.

Figure S3 presents the results of the simulation study for binary traits with a population prevalence of 1%. The study involved varying levels of genetic and residual correlations, with Cor($\mathbf{G_0}$, $\mathbf{G_E}$) values set at 0, 0.2, 0.4, 0.6 and 0.8, and Cor($\boldsymbol{\varepsilon}$, $\boldsymbol{\varepsilon}_\mathbf{E}$values set at 0, 0.2, 0.4, 0.6 and 0.8. In the discovery dataset, SNP effects were estimated using eq. (1) (GWAS) and eq. (2) (GWEIS). Model 1 has an inflated type 1 error rate especially at higher genetic correlation values (see Figure S3 a). When there is no genetic and residual correlation or no genetic and very low residual correlation (i.e. 0.2), type 1 error rate of Model 1 is controlled. Type 1 error rate of Model 2 possesses an unclear pattern, unlike Model 1. However, type 1 error rate of Model 3 is always controlled (at 5%). With respect to the statistical power, it is clear that Model 3 outperforms Models 1 and 2, where the power is higher in the presence of high GxE (see Figure S3 c and d). Power in Model 3 is relatively lower than the Models 1 and 2 in the presence of low GxE (see Figure S3 b), but the difference between the three models are negligible.

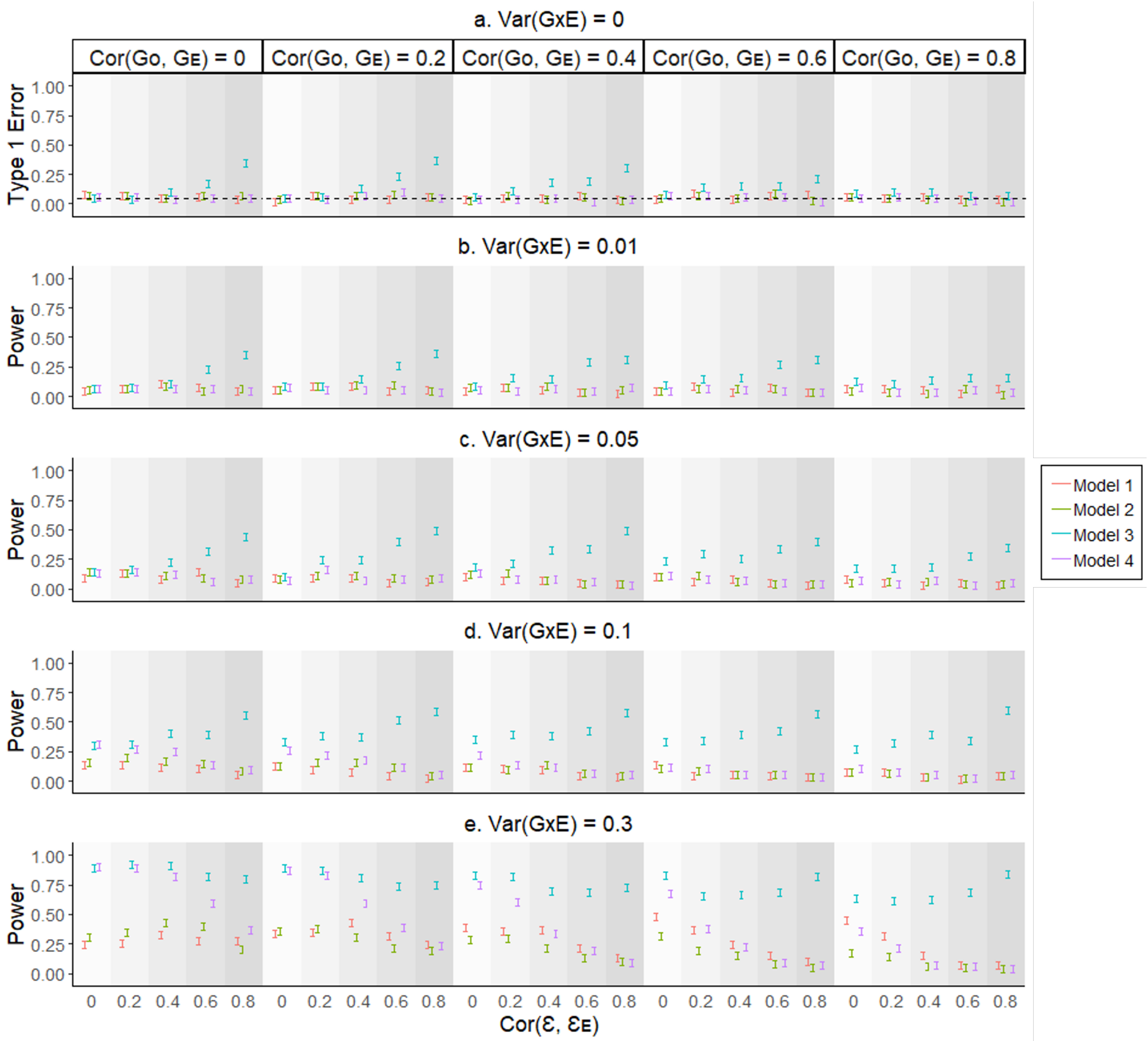

Figure S3: The type 1 error rate and power comparison of existing and proposed methods for binary traits with 1% population prevalence.

To assess the performance of our proposed method (Model 4) and existing methods (Models 1-3) on binary traits, we conducted simulation studies using various scenarios. We generated quantitative phenotypes and covariates using simulation models (eq. 7, 8, and 11) and converted them to binary phenotypes using a liability threshold of 1% population prevalence. We considered different levels of genetic and residual correlations (Cor($\mathbf{G_0}$, $\mathbf{G_E}$) = 0, 0.2, 0.4, 0.6 and 0.8), and Cor($\boldsymbol{\varepsilon}$, $\boldsymbol{\varepsilon_E}$) = 0, 0.2, 0.4, 0.6 and 0.8) and estimated SNP effects using GWAS (eq. 1) or GWEIS (eq. 2). We applied Models 1-4 to estimate the GxE component. The error bars in the figure represent the 95% confidence intervals for type 1 error rate and statistical power (vertical axes), and are based on averaging results from 200 simulated replicates.

Figure S4 presents the results of the simulation study for binary traits with a population prevalence of 50%. The study involved varying levels of genetic and residual correlations, with Cor($\mathbf{G_0}$, $\mathbf{G_E}$) values set at 0, 0.2, 0.4, 0.6 and 0.8, and Cor($\boldsymbol{\varepsilon}$, $\boldsymbol{\varepsilon_E}$values set at 0, 0.2, 0.4, 0.6 and 0.8. In the discovery dataset, SNP effects were estimated using eq. (1) (GWAS) and eq. (2) (GWEIS). Model 1 has an inflated type 1 error rate especially at higher genetic correlation values (see Figure S4 a). When there is no genetic and residual correlation or no genetic and very low residual correlation (i.e. 0.2), type 1 error rate of Model 1 is controlled. Type 1 error rate of Model 2 possesses an unclear pattern, unlike Model 1. However, type 1 error rate of Model 3 is always controlled (at 5%). With respect to the statistical power, it is clear that Model 3 outperforms Models 1 and 2, where the power is higher in the presence of high GxE (see Figure S4 c and d). Power in Model 3 is relatively lower than the Models 1 and 2 in the presence of low GxE (see Figure S4 b), but the difference between the three models are negligible.

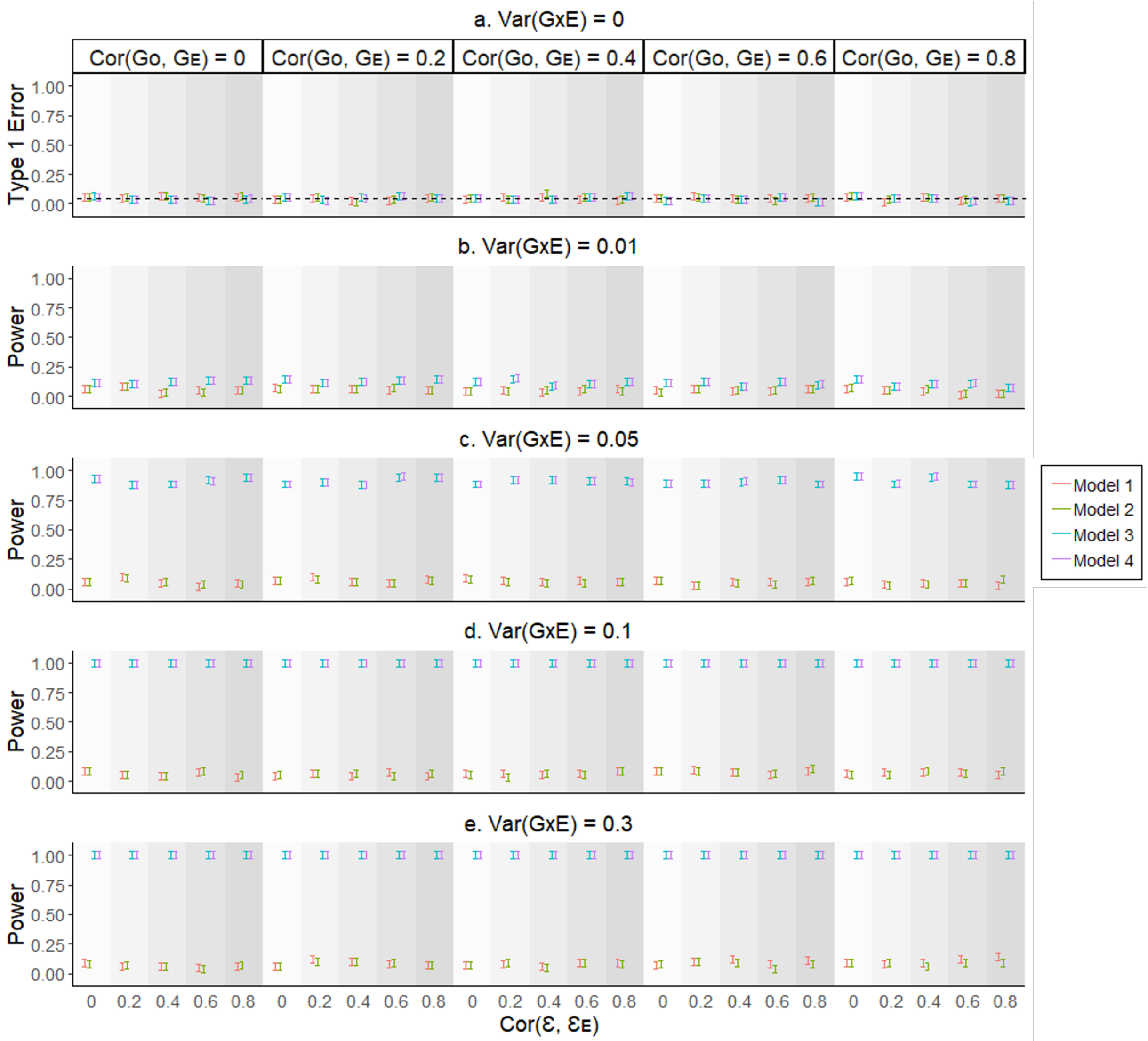

Figure S4: The type 1 error rate and power comparison of existing and proposed methods for binary traits with 50% population prevalence.

To assess the performance of our proposed method (Model 4) and existing methods (Models 1-3) on binary traits, we conducted simulation studies using various scenarios. We generated quantitative phenotypes and covariates using simulation models (eq. 7, 8, and 11) and converted them to binary phenotypes using a liability threshold of 50% population prevalence. We considered different levels of genetic and residual correlations (Cor($\mathbf{G_0}$, $\mathbf{G_E}$) = 0, 0.2, 0.4, 0.6 and 0.8), and Cor($\boldsymbol{\varepsilon}$, $\boldsymbol{\varepsilon}_\mathbf{E}$) = 0, 0.2, 0.4, 0.6 and 0.8) and estimated SNP effects using GWAS (eq. 1) or GWEIS (eq. 2). We applied Models 1-4 to estimate the GxE component. The error bars in the figure represent the 95% confidence intervals for type 1 error rate and statistical power (vertical axes), and are based on averaging results from 200 simulated replicates.

### Model Misspecification

#### Quantitative Traits

Figure S5 illustrates the findings of the simulation analysis for quantitative traits, when the model is misspecified. The simulation (true) models are given by eq. (9), (10) and (11). In the design, a range of values were selected for genetic and residual correlations. (Cor($\mathbf{G_0}$, $\mathbf{G_E}$) = 0, 0.2, 0.4, 0.6 and 0.8, and Cor($\boldsymbol{\varepsilon}$, $\boldsymbol{\varepsilon}_\mathbf{E}$) = 0, 0.2, 0.4, 0.6 and 0.8). We used permutation technique as a remedy to model misspecification. Despite the Var(RxE), in the absence of interaction term (i.e. Var(GxE) = 0) the permuted model of GxE PRS method (Model 4*) is controlled for type 1 error rate of 5% while other three models show inflation.

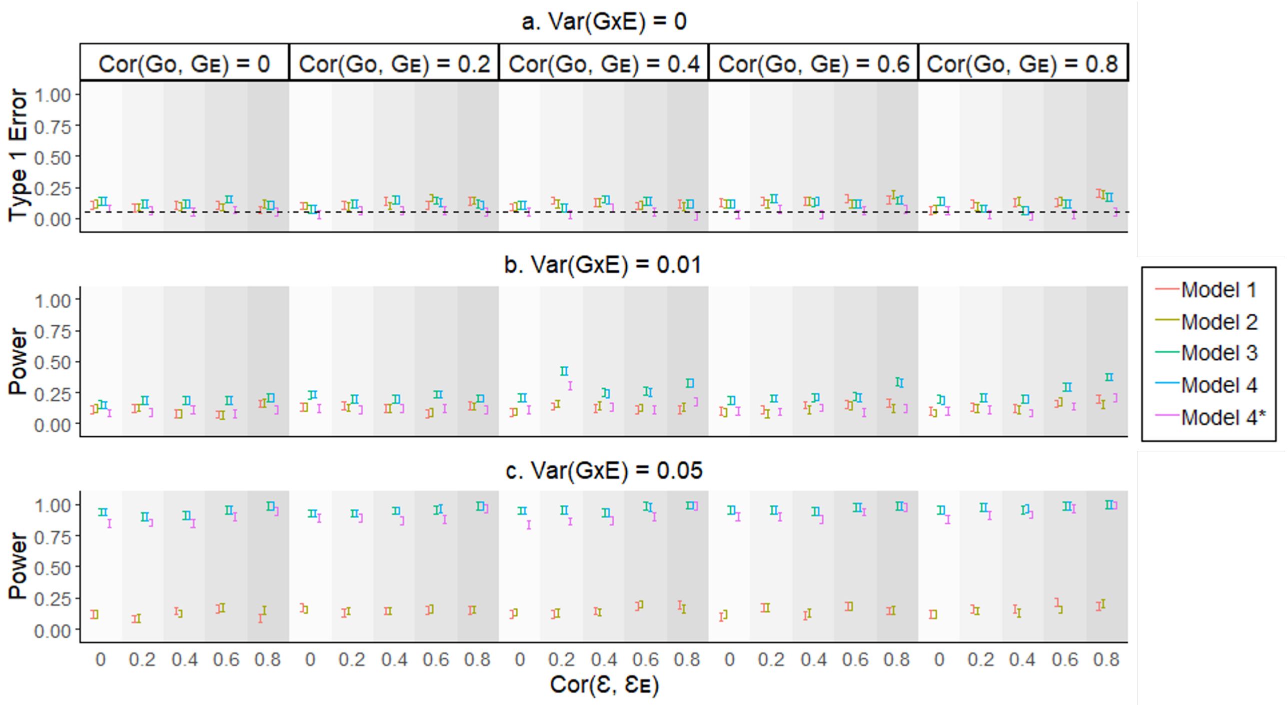

Figure S5: The type 1 error rate and statistical power of various GxE PRS models when using quantitative traits and Var(RxE) = 0.25.

We used simulation models (eq. 9, 10, and 11) to generate phenotypes and covariates. Different genetic and residual correlations (Cor($\mathbf{G_0}$, $\mathbf{G_E}$) = 0, 0.2, 0.4, 0.6 and 0.8), and Cor($\boldsymbol{\varepsilon}$, $\boldsymbol{\varepsilon_E}$) = 0, 0.2, 0.4, 0.6 and 0.8) were considered in various scenarios. Var(RxE) was set to 0.25. In the absence of GxE (eq. 9), we simulated the quantitative trait by adding the RxE component with the residual term. In the presence of GxE (eq. 10), we added the GxE component in addition to RxE and residual terms to simulate the quantitative trait. We applied Models 1 - 4 to estimate the GxE component, with SNP effects estimated from GWAS (eq. 1) or GWEIS (eq. 2). Model 4* is the permuted version of Model 4, obtained by permuting the $\hat{\mathbf{X}}_{\mathbf{gxe}}$ term of $\hat{\mathbf{X}}_{\mathbf{gxe}} \odot \mathbf{E}$ component 1000 times. The error bars show the 95% confidence intervals for type 1 error rate and statistical power (vertical axes), based on averaging results from 200 simulated replicates.

Figure S6 illustrates the findings of the simulation analysis for quantitative traits, when the model is misspecified. The simulation (true) models are given by eq. (9), (10) and (11). In the design, a range of values were selected for genetic and residual correlations. (Cor($\mathbf{G_0}$, $\mathbf{G_E}$) = 0, 0.2, 0.4, 0.6 and 0.8, and Cor($\boldsymbol{\varepsilon}$, $\boldsymbol{\varepsilon_E}$) = 0, 0.2, 0.4, 0.6 and 0.8). We used permutation technique as a remedy to model misspecification. Despite the Var(RxE), in the absence of interaction term (i.e. Var(GxE) = 0) the permuted model of GxE PRS method (Model 4*) is controlled for type 1 error rate of 5% while other three models show inflation.

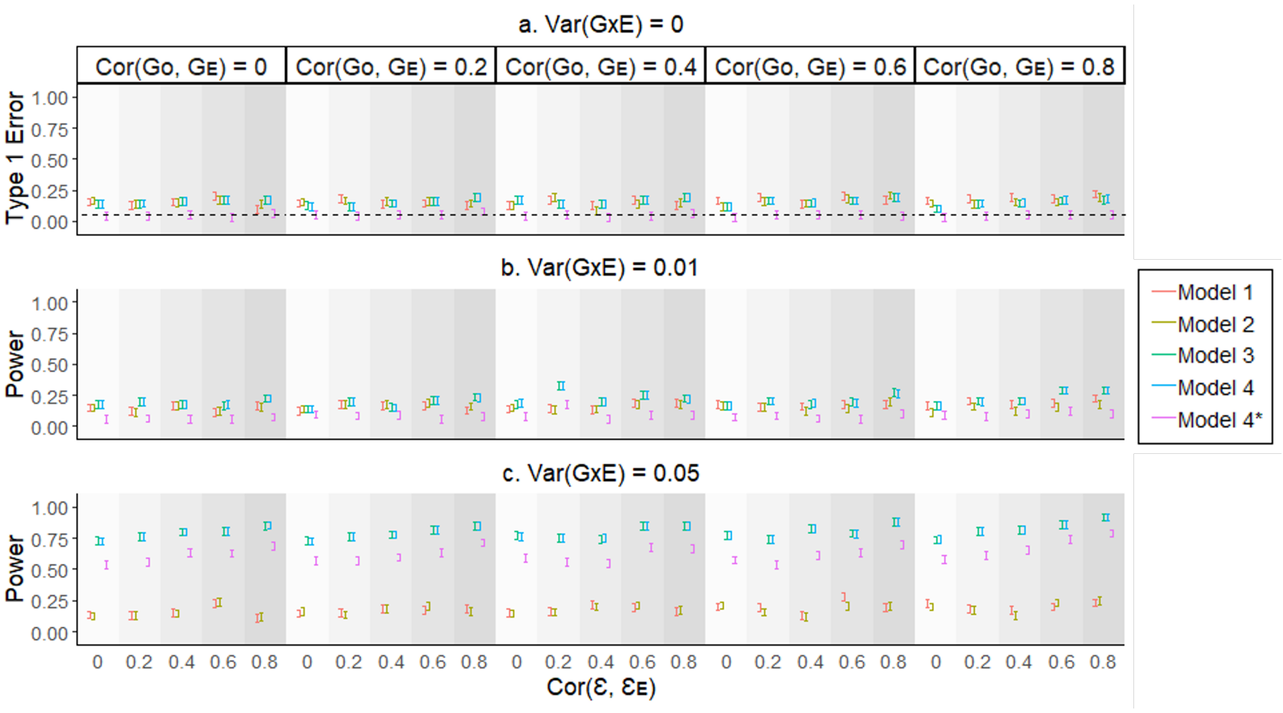

Figure S6: The type 1 error rate and statistical power of various GxE PRS models when using quantitative traits and Var(RxE) = 0.5.

We used simulation models (eq. 9, 10, and 11) to generate phenotypes and covariates. Different genetic and residual correlations (Cor($\mathbf{G_0}$, $\mathbf{G_E}$) = 0, 0.2, 0.4, 0.6 and 0.8), and Cor($\boldsymbol{\varepsilon}$, $\boldsymbol{\varepsilon_E}$) = 0, 0.2, 0.4, 0.6 and 0.8) were considered in various scenarios. Var(RxE) was set to 0.5. In the absence of GxE (eq. 9), we simulated the quantitative trait by adding the RxE component with the residual term. In the presence of GxE (eq. 10), we added the GxE component in addition to RxE and residual terms to simulate the quantitative trait. We applied Models 1 - 4 to estimate the GxE component, with SNP effects estimated from GWAS (eq. 1) or GWEIS (eq. 2). Model 4* is the permuted version of Model 4, obtained by permuting the $\hat{\mathbf{X}}_{\mathbf{gxe}}$ term of $\hat{\mathbf{X}}_{\mathbf{gxe}} \odot \mathbf{E}$ component 1000 times. The error bars show the 95% confidence intervals for type 1 error rate and statistical power (vertical axes), based on averaging results from 200 simulated replicates.

#### Binary Traits

Figure S7 illustrates the findings of the simulation analysis for binary traits, when the population prevalence is 10% and the model is misspecified. The simulation (true) models are given by eq. (9), (10) and (11). In the design, a range of values were selected for genetic and residual correlations. (Cor($\mathbf{G_0}$, $\mathbf{G_E}$) = 0, 0.2, 0.4, 0.6 and 0.8, and Cor($\boldsymbol{\varepsilon}$, $\boldsymbol{\varepsilon_E}$) = 0, 0.2, 0.4, 0.6 and 0.8). Despite the Var(RxE), in the absence of interaction term (i.e. Var(GxE) = 0) none of the models are controlled for type 1 error rate of 5%.

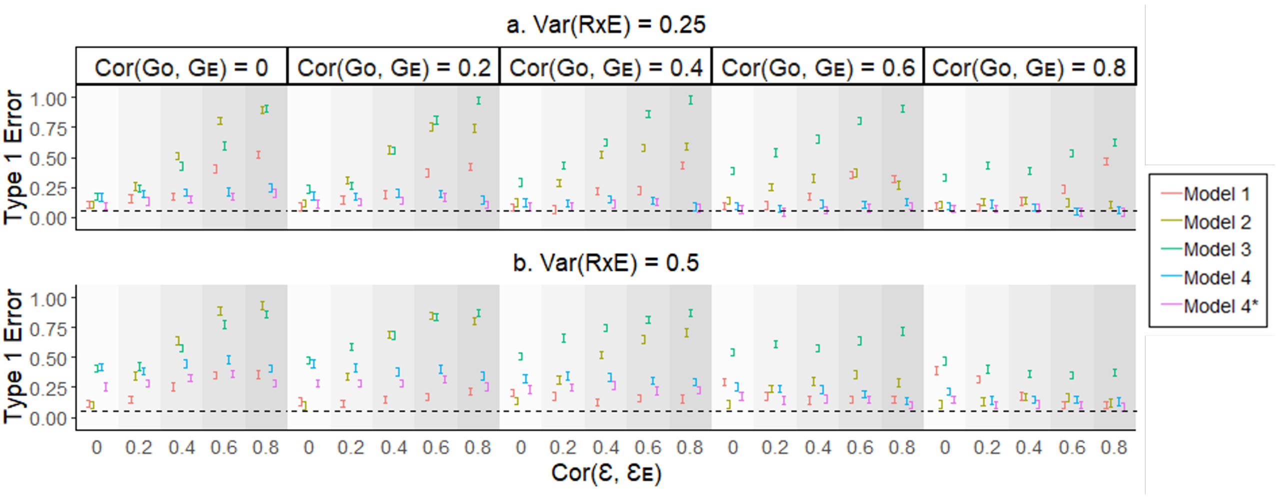

Figure S7: The type 1 error rate and statistical power of various GxE PRS models when using binary traits with Var(RxE) = 0.25 or 0.5.

We used simulation models (eq. 9, 10, and 11) to generate phenotypes and covariates. We used liability threshold of 10% population prevalence to simulate binary phenotypic outcomes. Different genetic and residual correlations (Cor($\mathbf{G_0}$, $\mathbf{G_E}$) = 0, 0.2, 0.4, 0.6 and 0.8), and Cor($\boldsymbol{\varepsilon}$, $\boldsymbol{\varepsilon_E}$) = 0, 0.2, 0.4, 0.6 and 0.8) were considered in various scenarios. In the absence of GxE (eq. 9), we simulated the quantitative trait by adding the RxE component with the residual term, and then converted to binary scale using liability threshold of 10% population prevalence. We applied Models 1 - 4 to estimate the GxE component, with SNP effects estimated from GWAS (eq. 1) or GWEIS (eq. 2). Model 4* is the permuted version of Model 4, obtained by permuting the $\hat{\mathbf{X}}_{\mathbf{gxe}}$ term of $\hat{\mathbf{X}}_{\mathbf{gxe}} \odot \mathbf{E}$ component 1000 times. The error bars show the 95% confidence intervals for type 1 error rate and statistical power (vertical axes), based on averaging results from 200 simulated replicates.

Figure S8 illustrates the findings of the simulation analysis for binary traits, when the population prevalence is 10% and the model is misspecified. The simulation (true) models are given by eq. (9), (10) and (11). In the design, a range of values were selected for genetic and residual correlations. (Cor($\mathbf{G_0}$, $\mathbf{G_E}$) = 0, 0.2, 0.4, 0.6 and 0.8, and Cor($\boldsymbol{\varepsilon}$, $\boldsymbol{\varepsilon_E}$) = 0, 0.2, 0.4, 0.6 and 0.8). When the model of the GxE PRS method is adjusted by incorporating the second order covariate term ($\mathbf{E}^2$) in both discovery and target models (i.e. Model 5), it is clear that the adjusted model is controlled for type 1 error of 5%, when Var(RxE) is 0.25. However, Model 5 does not show significantly higher power than the other models, but, it is clear that the power is increased at the presence of high GxE interaction (see Figure S8 b, c and d).

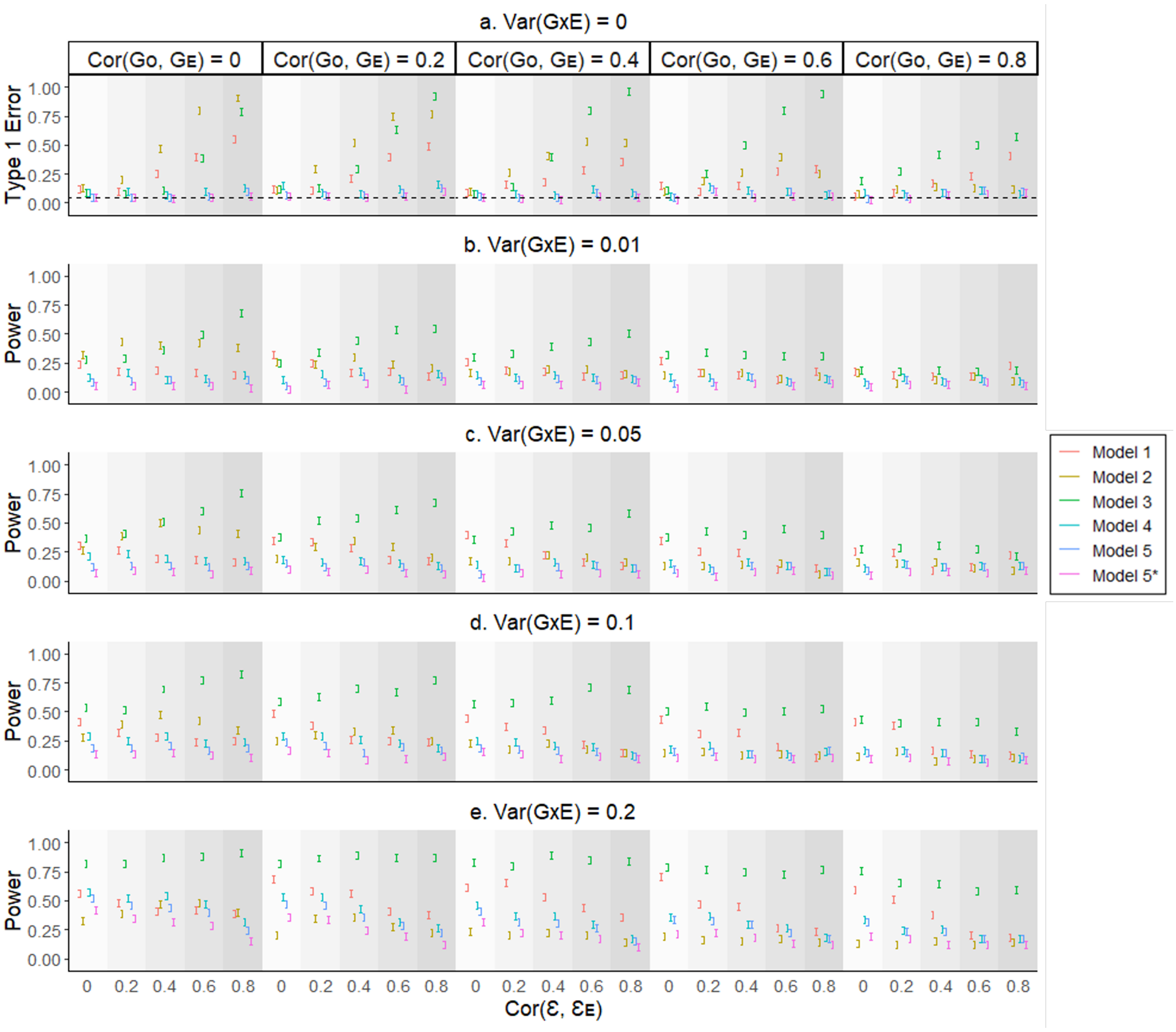

Figure S8: Type 1 error rate and power of different GxE PRS models when the phenotype is binary with 10% population prevalence and Var(RxE) = 0.25

To investigate the impact of model misspecification, we generated phenotypes and covariates with RxE effects using simulation models (eq. 9 and 10). The genetic and residual variances were set to fixed values of 0.4 and 0.5, respectively, for the main phenotypes. We used liability threshold of 10% population prevalence to simulate binary phenotypic outcomes. Different genetic and residual correlations (Cor($\mathbf{G_0}$, $\mathbf{G_E}$) = 0, 0.2, 0.4, 0.6 and 0.8), and Cor($\boldsymbol{\varepsilon}$, $\boldsymbol{\varepsilon_E}$) = 0, 0.2, 0.4, 0.6 and 0.8) were considered in various scenarios. Var(RxE) was set to 0.25. In the absence of GxE (eq. 9), we simulated the quantitative trait by adding the RxE component with the residual term, and then converted to binary scale using liability threshold of 10% population prevalence. In the presence of GxE (eq. 10), we added the GxE component in addition to RxE and residual terms to simulate the quantitative trait and converted to binary scale in a similar manner. We applied Models 1 - 5, with SNP effects estimated from GWAS (eq. (1)) or GWEIS (eq. (3)). Model 5* is the permuted version of Model 5, obtained by permuting the $\hat{\mathbf{X}}_{\mathbf{gxe}}$ term of $\hat{\mathbf{X}}_{\mathbf{gxe}} \odot \mathbf{E}$ component 1000 times. The error bars represent the 95% confidence intervals of the type 1 error rate and statistical power (vertical axes), based on averaging the results from 200 simulated replicates.

Figure S9 illustrates the findings of the simulation analysis for binary traits, when the population prevalence is 10% and the model is misspecified. The simulation (true) models are given by eq. (9), (10) and (11). In the design, a range of values were selected for genetic and residual correlations. (Cor($\mathbf{G_0}$, $\mathbf{G_E}$) = 0, 0.2, 0.4, 0.6 and 0.8, and Cor($\boldsymbol{\varepsilon}$, $\boldsymbol{\varepsilon_E}$) = 0, 0.2, 0.4, 0.6 and 0.8). When the model of the GxE PRS method is adjusted by incorporating the second order covariate term ($\mathbf{E}^2$) in both discovery and target models (i.e. Model 5), it is clear that the adjusted model is controlled for type 1 error of 5%, when Var(RxE) is 0.5. However, Model 5 does not show significantly higher power than the other models, but, it is clear that the power is increased at the presence of high GxE interaction (see Figure S9 b, c and d).

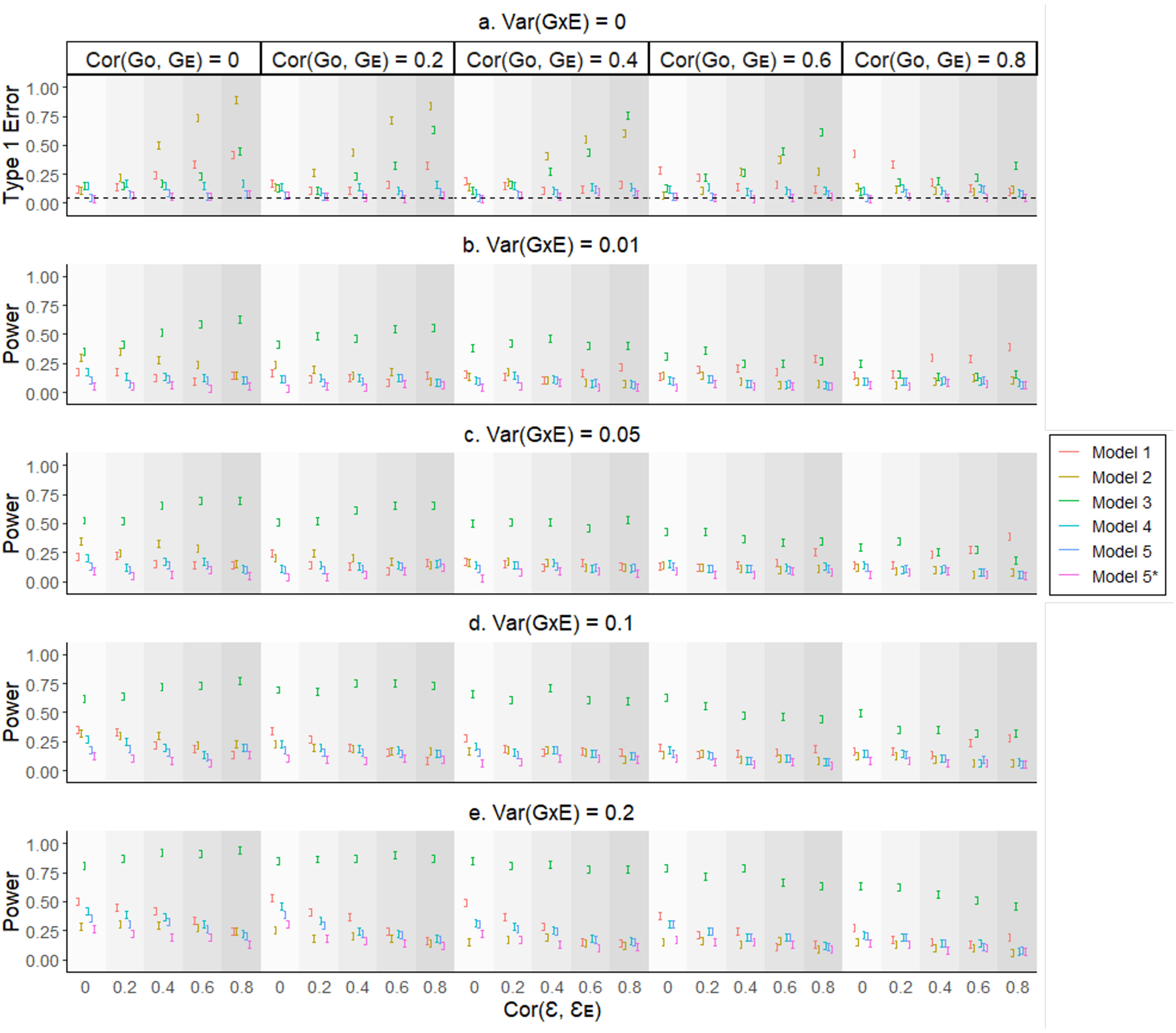

Figure S9: Type 1 error rate and power of different GxE PRS models when the phenotype is binary with 10% population prevalence and Var(RxE) = 0.5

To investigate the impact of model misspecification, we generated phenotypes and covariates with RxE effects using simulation models (eq. 9 and 10). The genetic and residual variances were set to fixed values of 0.4 and 0.5, respectively, for the main phenotypes. We used liability threshold of 10% population prevalence to simulate binary phenotypic outcomes. Different genetic and residual correlations (Cor($\mathbf{G_0}$, $\mathbf{G_E}$) = 0, 0.2, 0.4, 0.6 and 0.8), and Cor($\boldsymbol{\varepsilon}$, $\boldsymbol{\varepsilon_E}$) = 0, 0.2, 0.4, 0.6 and 0.8) were considered in various scenarios. Var(RxE) was set to 0.5. In the absence of GxE (eq. 9), we simulated the quantitative trait by adding the RxE component with the residual term, and then converted to binary scale using liability threshold of 10% population prevalence. In the presence of GxE (eq. 10), we added the GxE component in addition to RxE and residual terms to simulate the quantitative trait and converted to binary scale in a similar manner. We applied Models 1 - 5, with SNP effects estimated from GWAS (eq. (1)) or GWEIS (eq. (3)). Model 5* is the permuted version of Model 5, obtained by permuting the $\hat{\mathbf{X}}_{\mathbf{gxe}}$ term of $\hat{\mathbf{X}}_{\mathbf{gxe}} \odot \mathbf{E}$ component 1000 times. The error bars represent the 95% confidence intervals of the type 1 error rate and statistical power (vertical axes), based on averaging the results from 200 simulated replicates.

Figure S10 illustrates the findings of the simulation analysis for binary traits, when the population prevalence is 1% and the model is misspecified. The simulation (true) models are given by eq. (9), (10) and (11). In the design, a range of values were selected for genetic and residual correlations. (Cor($\mathbf{G_0}$, $\mathbf{G_E}$) = 0, 0.2, 0.4, 0.6 and 0.8, and Cor($\boldsymbol{\varepsilon}$, $\boldsymbol{\varepsilon_E}$) = 0, 0.2, 0.4, 0.6 and 0.8). When the model of the GxE PRS method is adjusted by incorporating the second order covariate term ($\mathbf{E}^2$) in both discovery and target models (i.e. Model 5), it is clear that the adjusted model is controlled for type 1 error of 5%, when Var(RxE) is 0.25. However, Model 5 does not show significantly higher power than the other models (see Figure S10 b, c and d).

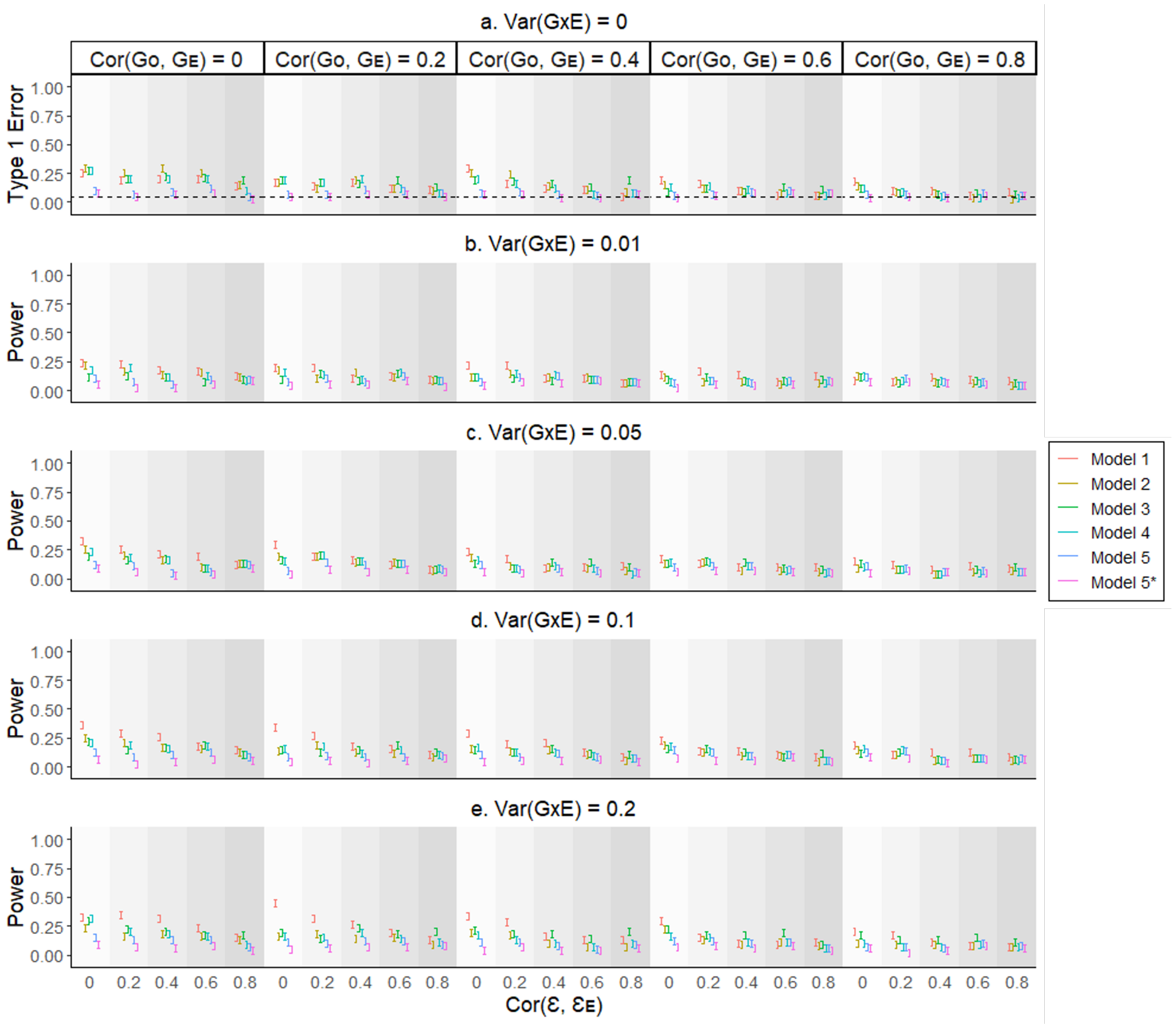

Figure S10: Type 1 error rate and power of different GxE PRS models when the phenotype is binary with 1% population prevalence and Var(RxE) = 0.25

To investigate the impact of model misspecification, we generated phenotypes and covariates with RxE effects using simulation models (eq. 9 and 10). The genetic and residual variances were set to fixed values of 0.4 and 0.5, respectively, for the main phenotypes. We used liability threshold of 1% population prevalence to simulate binary phenotypic outcomes. Different genetic and residual correlations (Cor($\mathbf{G_0}$, $\mathbf{G_E}$) = 0, 0.2, 0.4, 0.6 and 0.8), and Cor($\boldsymbol{\varepsilon}$, $\boldsymbol{\varepsilon_E}$) = 0, 0.2, 0.4, 0.6 and 0.8) were considered in various scenarios. Var(RxE) was set to 0.25. In the absence of GxE (eq. 9), we simulated the quantitative trait by adding the RxE component with the residual term, and then converted to binary scale using liability threshold of 1% population prevalence. In the presence of GxE (eq. 10), we added the GxE component in addition to RxE and residual terms to simulate the quantitative trait and converted to binary scale in a similar manner. We applied Models 1 - 5, with SNP effects estimated from GWAS (eq. (1)) or GWEIS (eq. (3)). Model 5* is the permuted version of Model 5, obtained by permuting the $\hat{\mathbf{X}}_{\mathbf{gxe}}$ term of $\hat{\mathbf{X}}_{\mathbf{gxe}} \odot \mathbf{E}$ component 1000 times. The error bars represent the 95% confidence intervals of the type 1 error rate and statistical power (vertical axes), based on averaging the results from 200 simulated replicates.

Figure S11 illustrates the findings of the simulation analysis for binary traits, when the population prevalence is 1% and the model is misspecified. The simulation (true) models are given by eq. (9), (10) and (11). In the design, a range of values were selected for genetic and residual correlations. (Cor($\mathbf{G_0}$, $\mathbf{G_E}$) = 0, 0.2, 0.4, 0.6 and 0.8, and Cor($\boldsymbol{\varepsilon}$, $\boldsymbol{\varepsilon_E}$) = 0, 0.2, 0.4, 0.6 and 0.8). When the model of the GxE PRS method is adjusted by incorporating the second order covariate term ($\mathbf{E}^2$) in both discovery and target models (i.e. Model 5), it is clear that the adjusted model is controlled for type 1 error of 5%, when Var(RxE) is 0.5. However, Model 5 does not show significantly higher power than the other models (see Figure S11 b, c and d).

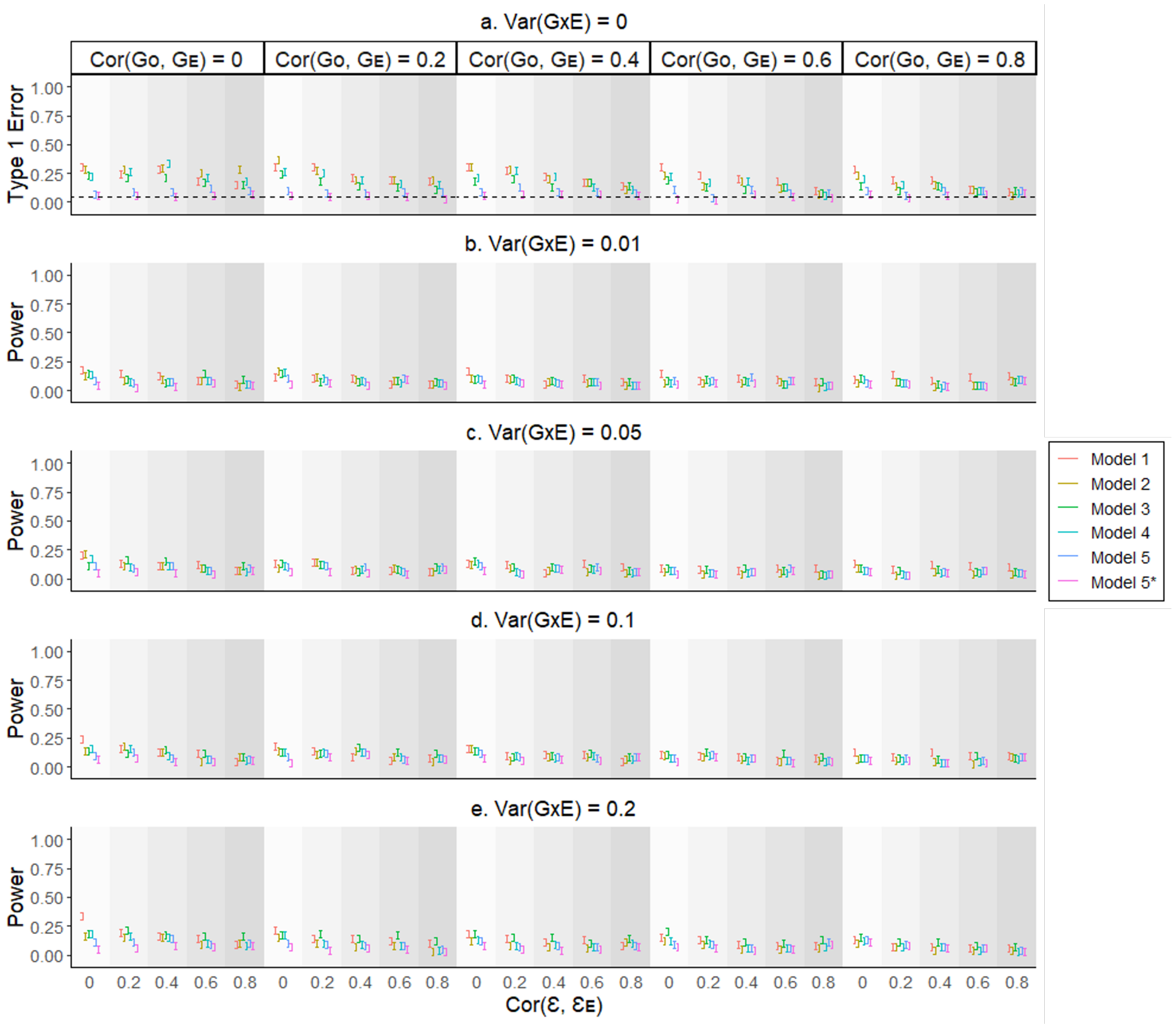

Figure S11: Type 1 error rate and power of different GxE PRS models when the phenotype is binary with 1% population prevalence and Var(RxE) = 0.5

To investigate the impact of model misspecification, we generated phenotypes and covariates with RxE effects using simulation models (eq. 9 and 10). The genetic and residual variances were set to fixed values of 0.4 and 0.5, respectively, for the main phenotypes. We used liability threshold of 1% population prevalence to simulate binary phenotypic outcomes. Different genetic and residual correlations (Cor($\mathbf{G_0}$, $\mathbf{G_E}$) = 0, 0.2, 0.4, 0.6 and 0.8), and Cor($\boldsymbol{\varepsilon}$, $\boldsymbol{\varepsilon}_\mathbf{E}$) = 0, 0.2, 0.4, 0.6 and 0.8) were considered in various scenarios. Var(RxE) was set to 0.5. In the absence of GxE (eq. 9), we simulated the quantitative trait by adding the RxE component with the residual term, and then converted to binary scale using liability threshold of 1% population prevalence. In the presence of GxE (eq. 10), we added the GxE component in addition to RxE and residual terms to simulate the quantitative trait and converted to binary scale in a similar manner. We applied Models 1 - 5, with SNP effects estimated from GWAS (eq. (1)) or GWEIS (eq. (3)). Model 5* is the permuted version of Model 5, obtained by permuting the $\hat{\mathbf{X}}_{\mathbf{gxe}}$ term of $\hat{\mathbf{X}}_{\mathbf{gxe}} \odot \mathbf{E}$ component 1000 times. The error bars represent the 95% confidence intervals of the type 1 error rate and statistical power (vertical axes), based on averaging the results from 200 simulated replicates.

Figure S12 illustrates the findings of the simulation analysis for binary traits, when the population prevalence is 50% and the model is misspecified. The simulation (true) models are given by eq. (9), (10) and (11). In the design, a range of values were selected for genetic and residual correlations. (Cor($\mathbf{G_0}$, $\mathbf{G_E}$) = 0, 0.2, 0.4, 0.6 and 0.8, and Cor($\boldsymbol{\varepsilon}$, $\boldsymbol{\varepsilon}_\mathbf{E}$) = 0, 0.2, 0.4, 0.6 and 0.8). When the model of the GxE PRS method is adjusted by incorporating the second order covariate term ($\mathbf{E}^2$) in both discovery and target models (i.e. Model 5), it is clear that all the models are controlled for type 1 error of 5%, when Var(RxE) is 0.25. However, Model 5 does not show significantly higher power than the other models in lower GxE (see Figure S12 b). Models 3, 4 and 5 show significantly higher power than Models 1 and 2 in higher GxE (see Figure S12 c and d).

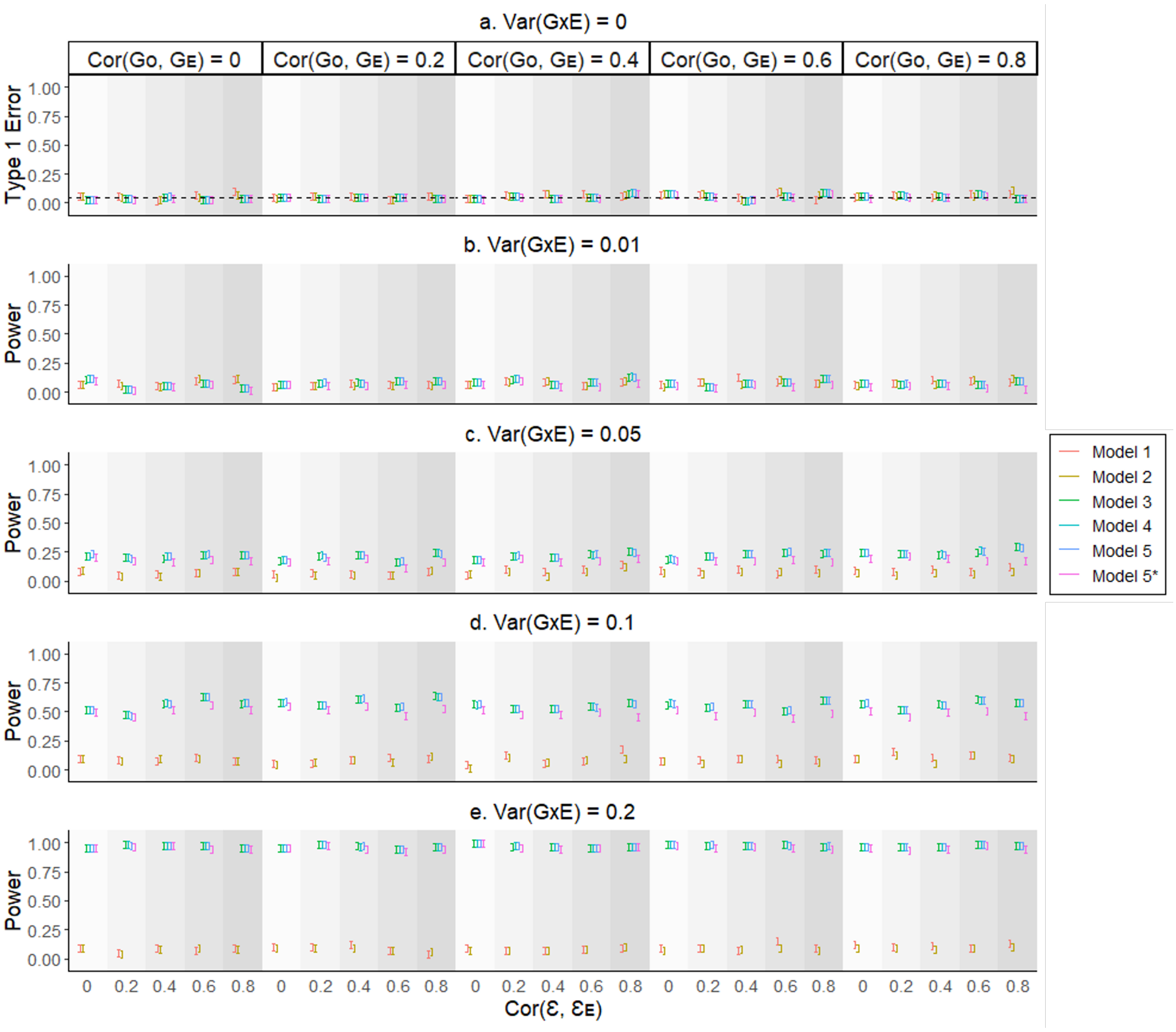

Figure S12: Type 1 error rate and power of different GxE PRS models when the phenotype is binary with 50% population prevalence and Var(RxE) = 0.25

To investigate the impact of model misspecification, we generated phenotypes and covariates with RxE effects using simulation models (eq. 9 and 10). The genetic and residual variances were set to fixed values of 0.4 and 0.5, respectively, for the main phenotypes. We used liability threshold of 50% population prevalence to simulate binary phenotypic outcomes. Different genetic and residual correlations (Cor($\mathbf{G_0}$, $\mathbf{G_E}$) = 0, 0.2, 0.4, 0.6 and 0.8), and Cor($\boldsymbol{\varepsilon}$, $\boldsymbol{\varepsilon_E}$) = 0, 0.2, 0.4, 0.6 and 0.8) were considered in various scenarios. Var(RxE) was set to 0.25. In the absence of GxE (eq. 9), we simulated the quantitative trait by adding the RxE component with the residual term, and then converted to binary scale using liability threshold of 50% population prevalence. In the presence of GxE (eq. 10), we added the GxE component in addition to RxE and residual terms to simulate the quantitative trait and converted to binary scale in a similar manner. We applied Models 1 - 5, with SNP effects estimated from GWAS (eq. (1)) or GWEIS (eq. (3)). Model 5* is the permuted version of Model 5, obtained by permuting the $\hat{\mathbf{X}}_{\mathbf{gxe}}$ term of $\hat{\mathbf{X}}_{\mathbf{gxe}} \odot \mathbf{E}$ component 1000 times. The error bars represent the 95% confidence intervals of the type 1 error rate and statistical power (vertical axes), based on averaging the results from 200 simulated replicates.

Figure S13 illustrates the findings of the simulation analysis for binary traits, when the population prevalence is 50% and the model is misspecified. The simulation (true) models are given by eq. (9), (10) and (11). In the design, a range of values were selected for genetic and residual correlations. (Cor($\mathbf{G_0}$, $\mathbf{G_E}$) = 0, 0.2, 0.4, 0.6 and 0.8, and Cor($\boldsymbol{\varepsilon}$, $\boldsymbol{\varepsilon_E}$) = 0, 0.2, 0.4, 0.6 and 0.8). When the model of the GxE PRS method is adjusted by incorporating the second order covariate term ($\mathbf{E}^2$) in both discovery and target models (i.e. Model 5), it is clear that all the models are controlled for type 1 error of 5%, when Var(RxE) is 0.25. However, Model 5 does not show significantly higher power than the other models in lower GxE (see Figure S13 b). Models 3, 4 and 5 show significantly higher power than Models 1 and 2 in higher GxE (see Figure S13 c and d).

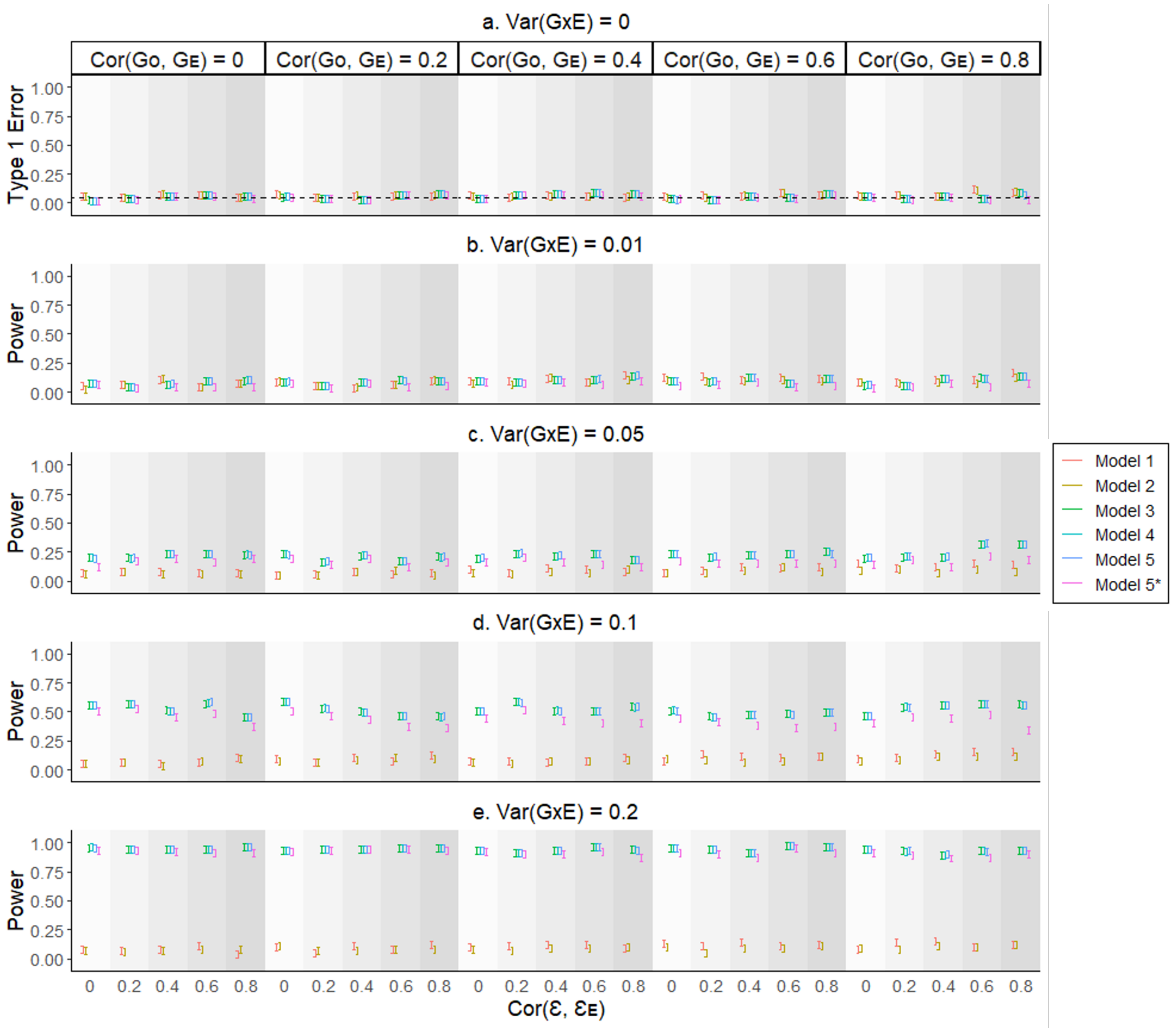

Figure S13: Type 1 error rate and power of different GxE PRS models when the phenotype is binary with 50% population prevalence and Var(RxE) = 0.5

To investigate the impact of model misspecification, we generated phenotypes and covariates with RxE effects using simulation models (eq. 9 and 10). The genetic and residual variances were set to fixed values of 0.4 and 0.5, respectively, for the main phenotypes. We used liability threshold of 50% population prevalence to simulate binary phenotypic outcomes. Different genetic and residual correlations (Cor($\mathbf{G_0}$, $\mathbf{G_E}$) = 0, 0.2, 0.4, 0.6 and 0.8), and Cor($\boldsymbol{\varepsilon}$, $\boldsymbol{\varepsilon_E}$) = 0, 0.2, 0.4, 0.6 and 0.8) were considered in various scenarios. Var(RxE) was set to 0.5. In the absence of GxE (eq. 9), we simulated the quantitative trait by adding the RxE component with the residual term, and then converted to binary scale using liability threshold of 50% population prevalence. In the presence of GxE (eq. 10), we added the GxE component in addition to RxE and residual terms to simulate the quantitative trait and converted to binary scale in a similar manner. We applied Models 1 - 5, with SNP effects estimated from GWAS (eq. (1)) or GWEIS (eq. (3)). Model 5* is the permuted version of Model 5, obtained by permuting the $\mathbf{\hat{X}_{gxe}}$ term of $\mathbf{\hat{X}_{gxe}} \odot \mathbf{E}$ component 1000 times. The error bars represent the 95% confidence intervals of the type 1 error rate and statistical power (vertical axes), based on averaging the results from 200 simulated replicates.

### Real Data

Table S1 contains the details of quantitative scale variables extracted from the UK BioBank which were considered as quantitative outcomes and/or covariates in this article. Anthropometric measures were obtained through physical examination and health behaviour measures were collected from participants via researcher-administered surveys upon entry into the UK Biobank study. Serum marker measures were derived from analysis of participant blood samples taken at baseline.

Table S1: Quantitative variable details

| UK biobank Field ID | Variable | Abbreviation |
|---|---|---|
| 1289 | Cooked vegetable intake | HD |
| 1299 | Salad/raw vegetable intake | |
| 1309 | Fresh fruit intake | |
| 1329 | Oily fish intake | |
| 1339 | Non-oily fish intake | |
| 1379 | Lamb/mutton intake | |
| 1349 | Processed meat intake | |
| 1369 | Beef intake | |
| 1389 | Pork intake | |
| 48 | Waist circumference | WHR |
| 49 | Hip circumference | |
| 21001 | Body mass index | BMI |
| 1558 | Alcohol intake frequency | ALC |
| 22040 | Summed MET minutes per week for all activity | PA |
| 30760 | HDL cholesterol | HDL |
| 30020 | Haemoglobin concentration | HGB |
| 30780 | LDL direct | LDL |

We considered 1) fruit and vegetable intake, 2) total fish intake (oily and non-oily), 3) processed meat intake and 4) red meat intake (beef, pork and lamb/mutton) as an index of healthy diet (HD) following the American Heart Association Guidelines[34].

1. The amount of total fruit and vegetable intake per day was obtained by combining the amount of total fruit servings (piece of fruit = 1 serving) and vegetable servings (3 tablespoons = 1 serving) consumed. Scores of individuals who had 4.5 or more servings a day were converted to 1, and the remaining scores were converted to 0.
2. Total fish intake was obtained by combining the data of oily and non-oily fish intake after recoding weekly frequency (0 = never, 0.5 = less than once week, 1 = once a week, 3 = two to four times a week, 5.5 = five to six times a week, and 7 = once or more daily). Scores of individuals who ate 2 or more times per week were converted to 1, and remaining intake frequencies were converted to 0.
3. Total processed meat intake a week was recorded as weekly intake (0 = never, 0.5 = less than once week, 1 = once a week, 3 = two to four times a week, 5.5 = five to six times a week, and 7 = once or more daily). Scores of individuals who consumed processed meat 2 or fewer times per week were converted to 1, and the remaining intake frequencies were converted to 0.
4. Total red meat intake was obtained from combining the data for beef, lamb/mutton and pork intake based on weekly intake (0 = never, 0.5 = less than once week, 1 = once a week, 3 = two to four times a week, 5.5 = five to six times a week, and 7 = once or more daily). The records of individuals who had 5 or fewer times per week were converted to 1, and the remaining intake frequencies were converted to 0.

The overall dietary score was derived by totaling the values for separate dietary components described above and scores ranged from a minimum of 0 point to a maximum of 4 point for each participant. In summary, a score close to 4 were represented as indicators of healthier diet habits than a score close to zero.

Waist-to-hip ratio (WHR) was computed by obtaining the ratio between waist circumference and hip circumference of corresponding individuals.

We re-coded the original alcohol intake frequency (ALC) so that increasing magnitude represented greater frequency of consumption (0 = never, 1 = special occasions only, 2 = one to three times a month, 3 = once or twice a week, 4 = three or four times a week and 5 = daily)[35].

Table S2 contains the details of binary scale variables used to describe disease status classification considering prevalent cases . We used ICD10 codes from UK biobank to capture diabetes (DIAB), hypertension (HYP) and coronary artery disease (CAD) cases.

Table S2: Binary variable details

| UK biobank Field ID | Variable | Remark |
|---|---|---|
| 41202 | Diagnoses - main ICD10 | Main Diagnoses |
| 41270 | Diagnosis - ICD10 | Summary Diagnoses |
| **Disease Code** | **Variable** | **Abbreviation** |
| E10 | Insulin-dependent diabetes mellitus | DIAB |
| E11 | Non-insulin-dependent diabetes mellitus | |
| E13 | Other specified diabetes mellitus | |
| E14 | Unspecified diabetes mellitus | |
| I10 | Essential (primary) hypertension | HYP |
| I11 | Hypertensive heart disease | |
| I12 | Hypertensive renal disease | |
| I13 | Hypertensive heart and renal disease | |
| I15 | Secondary hypertension | |
| I22 | Subsequent myocardial infarction | CAD |
| I23 | Certain current complications following acute myocardial infarction | |
| I24 | Other acute ischaemic heart diseases | |
| I25 | Chronic ischaemic heart disease | |

Table S3 contains the details of all the other variables used in this study.

Table S3: Other variable details

| UK biobank Field ID | Variable | Remark |
|---|---|---|
| 31 | Sex | Used to adjust phenotypes |
| 22189 | Townsend deprivation index at recruitment | |
| 21022 | Age at recruitment | |
| 6138 | Qualifications | |
| 22009 | Genetic principal components | |

Note that, the variable 'qualification' was based on the qualification questionnaire. We followed the International Standard Classification of Education[27] for converting the baseline qualification information to the education in years for each individual. We used the first 10 genetic principal components in our study.

#### Complete Real Data Analysis Results

Table S4: Analysis of BMI/ALC

| Model | Component | Estimate | Standard error | P value | R squared | Adjusted R squared |
|---|---|---|---|---|---|---|
| 1 | PRS_trd | 2.5937E-01 | 3.9407E-03 | <2E-16 | 6.7272E-02 | 1.1050E-01 |
| | PRS_trd x E | 5.1196E-02 | 4.5775E-03 | <2E-16 | 2.6211E-03 | |
| | E | 8.0568E-02 | 4.6742E-03 | <2E-16 | 6.4913E-03 | |
| 2 | PRS_add | 2.5799E-01 | 3.9403E-03 | <2E-16 | 6.6561E-02 | 1.0980E-01 |
| | PRS_add x E | 4.9088E-02 | 4.4494E-03 | <2E-16 | 2.4096E-03 | |
| | E | 8.6533E-02 | 4.5484E-03 | <2E-16 | 7.4879E-03 | |
| 3 | PRS_add | 2.5810E-01 | 3.9429E-03 | <2E-16 | 6.6615E-02 | 1.0860E-01 |
| | PRS_gxe x E | 2.7276E-02 | 4.0703E-03 | 2.0900E-11 | 7.4399E-04 | |
| | E | 1.0237E-01 | 4.1847E-03 | <2E-16 | 1.0479E-02 | |
| 4 | PRS_add | 2.6280E-01 | 4.2472E-03 | <2E-16 | 6.9065E-02 | 1.0870E-01 |
| | PRS_gxe | -1.2684E-02 | 4.2600E-03 | 2.9070E-03 | 1.6089E-04 | |
| | PRS_gxe x E | 2.7322E-02 | 4.0700E-03 | 1.9200E-11 | 7.4649E-04 | |
| | E | 1.0237E-01 | 4.1844E-03 | <2E-16 | 1.0479E-02 | |
| 4* | PRS_add | NA | NA | NA | NA | NA |
| | PRS_gxe | NA | NA | NA | NA | |
| | PRS_gxe x E | NA | NA | 0.0000E+00 | NA | |
| | E | NA | NA | NA | NA | |

Table S5: Analysis of BMI/HD

| Model | Component | Estimate | Standard error | P value | R squared | Adjusted R squared |
|---|---|---|---|---|---|---|
| 1 | PRS_trd | 2.6340E-01 | 3.9940E-03 | <2E-16 | 6.9377E-02 | 1.0160E-01 |
| | PRS_trd x E | -9.9627E-03 | 4.6526E-03 | 3.2250E-02 | 9.9255E-05 | |
| | E | -5.9869E-02 | 4.7607E-03 | <2E-16 | 3.5843E-03 | |
| 2 | PRS_add | 2.6118E-01 | 3.9964E-03 | <2E-16 | 6.8217E-02 | 1.0040E-01 |
| | PRS_add x E | -9.7895E-03 | 4.5523E-03 | 3.1500E-02 | 9.5834E-05 | |
| | E | -6.0983E-02 | 4.6626E-03 | <2E-16 | 3.7190E-03 | |
| 3 | PRS_add | 2.6127E-01 | 3.9963E-03 | <2E-16 | 6.8262E-02 | 1.0050E-01 |
| | PRS_gxe x E | 1.2549E-02 | 4.1434E-03 | 2.4600E-03 | 1.5747E-04 | |
| | E | -6.2308E-02 | 4.2632E-03 | <2E-16 | 3.8823E-03 | |
| 4 | PRS_add | 2.6142E-01 | 3.9967E-03 | <2E-16 | 6.8340E-02 | 1.0060E-01 |
| | PRS_gxe | -8.8305E-03 | 4.0102E-03 | 2.7670E-02 | 7.7978E-05 | |
| | PRS_gxe x E | 1.2499E-02 | 4.1433E-03 | 2.5600E-03 | 1.5623E-04 | |
| | E | -6.2381E-02 | 4.2632E-03 | <2E-16 | 3.8914E-03 | |
| 4* | PRS_add | NA | NA | NA | NA | NA |
| | PRS_gxe | NA | NA | NA | NA | |
| | PRS_gxe x E | NA | NA | 4.0000E-03 | NA | |
| | E | NA | NA | NA | NA | |

Table S6: Analysis of BMI/PA

| Model | Component | Estimate | Standard error | P value | R squared | Adjusted R squared |
|---|---|---|---|---|---|---|
| 1 | PRS_trd | 1.2437E+00 | 2.0252E-02 | <2E-16 | 1.5467E+00 | 1.1170E-01 |
| | PRS_trd x E | -1.5373E-01 | 2.3666E-02 | 8.3200E-11 | 2.3634E-02 | |
| | E | -4.0200E-01 | 2.3786E-02 | <2E-16 | 1.6161E-01 | |
| 2 | PRS_add | 1.2040E+00 | 2.0290E-02 | <2E-16 | 1.4496E+00 | 1.0720E-01 |
| | PRS_add x E | -1.2340E-01 | 2.1210E-02 | 5.9800E-09 | 1.5228E-02 | |
| | E | -4.5280E-01 | 2.1340E-02 | <2E-16 | 2.0503E-01 | |
| 3 | PRS_add | 1.2049E+00 | 2.0296E-02 | <2E-16 | 1.4518E+00 | 1.0670E-01 |
| | PRS_gxe x E | 5.5454E-02 | 2.8314E-02 | 5.0200E-02 | 3.0751E-03 | |
| | E | -4.5069E-01 | 2.8422E-02 | <2E-16 | 2.0312E-01 | |
| 4 | PRS_add | 1.2150E+00 | 2.0839E-02 | <2E-16 | 1.4761E+00 | 1.0670E-01 |
| | PRS_gxe | 4.4279E-02 | 2.0799E-02 | 3.3300E-02 | 1.9606E-03 | |
| | PRS_gxe x E | 5.5599E-02 | 2.8313E-02 | 4.9600E-02 | 3.0912E-03 | |
| | E | -4.5131E-01 | 2.8423E-02 | <2E-16 | 2.0368E-01 | |
| 4* | PRS_add | NA | NA | NA | NA | NA |
| | PRS_gxe | NA | NA | NA | NA | |
| | PRS_gxe x E | NA | NA | 5.7000E-02 | NA | |
| | E | NA | NA | NA | NA | |

Table S7: Analysis of LDL/ALC

| Model | Component | Estimate | Standard error | P value | R squared | Adjusted R squared |
|---|---|---|---|---|---|---|
| 1 | PRS_trd | 1.7032E-01 | 4.1742E-03 | <2E-16 | 2.9008E-02 | 4.2120E-02 |
| | PRS_trd x E | -2.9745E-03 | 5.0849E-03 | 5.5858E-01 | 8.8477E-06 | |
| | E | -2.9368E-02 | 5.1942E-03 | 1.5800E-08 | 8.6246E-04 | |
| 2 | PRS_add | 1.7033E-01 | 4.1743E-03 | <2E-16 | 2.9013E-02 | 4.2120E-02 |
| | PRS_add x E | -3.1988E-03 | 5.0488E-03 | 5.2636E-01 | 1.0232E-05 | |
| | E | -2.9940E-02 | 5.1586E-03 | 6.5200E-09 | 8.9640E-04 | |
| 3 | PRS_add | 1.7035E-01 | 4.1742E-03 | <2E-16 | 2.9018E-02 | 4.2120E-02 |
| | PRS_gxe x E | 2.5801E-03 | 4.5973E-03 | 5.7465E-01 | 6.6569E-06 | |
| | E | -2.9209E-02 | 4.7069E-03 | 5.4900E-10 | 8.5317E-04 | |
| 4 | PRS_add | 1.7081E-01 | 4.1775E-03 | <2E-16 | 2.9177E-02 | 4.2230E-02 |
| | PRS_gxe | -1.1432E-02 | 4.1855E-03 | 6.3100E-03 | 1.3068E-04 | |
| | PRS_gxe x E | 2.5745E-03 | 4.5970E-03 | 5.7546E-01 | 6.6281E-06 | |
| | E | -2.9285E-02 | 4.7067E-03 | 4.9500E-10 | 8.5759E-04 | |
| 4* | PRS_add | NA | NA | NA | NA | NA |
| | PRS_gxe | NA | NA | NA | NA | |
| | PRS_gxe x E | NA | NA | 5.7500E-01 | NA | |
| | E | NA | NA | NA | NA | |

Table S8: Analysis of LDL/HD

| Model | Component | Estimate | Standard error | P value | R squared | Adjusted R squared |
|---|---|---|---|---|---|---|
| 1 | PRS_trd | 1.6989E-01 | 4.2235E-03 | <2E-16 | 2.8861E-02 | 4.1630E-02 |
| | PRS_trd x E | 3.5915E-03 | 5.1067E-03 | 4.8188E-01 | 1.2899E-05 | |
| | E | -1.9504E-02 | 5.2149E-03 | 1.8400E-04 | 3.8041E-04 | |
| 2 | PRS_add | 1.6811E-01 | 4.2252E-03 | <2E-16 | 2.8262E-02 | 4.1030E-02 |
| | PRS_add x E | 4.4924E-03 | 5.2417E-03 | 3.9142E-01 | 2.0182E-05 | |
| | E | -1.9143E-02 | 5.3474E-03 | 3.4400E-04 | 3.6645E-04 | |
| 3 | PRS_add | 1.6808E-01 | 4.2252E-03 | <2E-16 | 2.8252E-02 | 4.1030E-02 |
| | PRS_gxe x E | -4.1325E-03 | 4.6795E-03 | 3.7718E-01 | 1.7078E-05 | |
| | E | -2.3576E-02 | 4.7984E-03 | 8.9800E-07 | 5.5582E-04 | |
| 4 | PRS_add | 1.6805E-01 | 4.2259E-03 | <2E-16 | 2.8241E-02 | 4.1010E-02 |
| | PRS_gxe | 1.8698E-03 | 4.2331E-03 | 6.5871E-01 | 3.4962E-06 | |
| | PRS_gxe x E | -4.1347E-03 | 4.6795E-03 | 3.7693E-01 | 1.7096E-05 | |
| | E | -2.3579E-02 | 4.7985E-03 | 8.9500E-07 | 5.5598E-04 | |
| 4* | PRS_add | NA | NA | NA | NA | NA |
| | PRS_gxe | NA | NA | NA | NA | |
| | PRS_gxe x E | NA | NA | 3.5800E-01 | NA | |
| | E | NA | NA | NA | NA | |

Table S9: Analysis of LDL/PA

| Model | Component | Estimate | Standard error | P value | R squared | Adjusted R squared |
|---|---|---|---|---|---|---|
| 1 | PRS_trd | 1.6938E-01 | 4.6005E-03 | <2E-16 | 2.8689E-02 | 3.9740E-02 |
| | PRS_trd x E | -7.0435E-03 | 5.5848E-03 | 2.0720E-01 | 4.9611E-05 | |
| | E | 9.9883E-03 | 5.6195E-03 | 7.5500E-02 | 9.9766E-05 | |
| 2 | PRS_add | 1.6124E-01 | 4.6122E-03 | <2E-16 | 2.5997E-02 | 3.6960E-02 |
| | PRS_add x E | -7.5874E-03 | 5.1418E-03 | 1.4010E-01 | 5.7569E-05 | |
| | E | 1.0761E-02 | 5.1861E-03 | 3.8000E-02 | 1.1581E-04 | |
| 3 | PRS_add | 1.6119E-01 | 4.6122E-03 | <2E-16 | 2.5981E-02 | 3.6920E-02 |
| | PRS_gxe x E | 2.6418E-03 | 4.6134E-03 | 5.6690E-01 | 6.9791E-06 | |
| | E | 1.4186E-02 | 4.6399E-03 | 2.2300E-03 | 2.0125E-04 | |
| 4 | PRS_add | 1.6121E-01 | 4.6125E-03 | <2E-16 | 2.5988E-02 | 3.6900E-02 |
| | PRS_gxe | 2.2549E-03 | 4.6109E-03 | 6.2481E-01 | 5.0846E-06 | |
| | PRS_gxe x E | 2.6159E-03 | 4.6138E-03 | 5.7074E-01 | 6.8429E-06 | |
| | E | 1.4186E-02 | 4.6399E-03 | 2.2300E-03 | 2.0124E-04 | |
| 4* | PRS_add | NA | NA | NA | NA | NA |
| | PRS_gxe | NA | NA | NA | NA | |
| | PRS_gxe x E | NA | NA | 5.6300E-01 | NA | |
| | E | NA | NA | NA | NA | |

Table S10: Analysis of WHR/ALC

| Model | Component | Estimate | Standard error | P value | R squared | Adjusted R squared |
|---|---|---|---|---|---|---|
| 1 | PRS_trd | 1.6591E-01 | 2.9786E-03 | <2E-16 | 2.7526E-02 | 4.9150E-01 |
| | PRS_trd x E | 2.2891E-02 | 3.2545E-03 | 2.0300E-12 | 5.2401E-04 | |
| | E | 3.8260E-02 | 3.3391E-03 | <2E-16 | 1.4638E-03 | |
| 2 | PRS_add | 1.6558E-01 | 2.9784E-03 | <2E-16 | 2.7417E-02 | 4.9140E-01 |
| | PRS_add x E | 2.2994E-02 | 3.3451E-03 | 6.3100E-12 | 5.2871E-04 | |
| | E | 4.0276E-02 | 3.4273E-03 | <2E-16 | 1.6221E-03 | |
| 3 | PRS_add | 1.6578E-01 | 2.9792E-03 | <2E-16 | 2.7484E-02 | 4.9110E-01 |
| | PRS_gxe x E | 8.6652E-03 | 2.9707E-03 | 3.5400E-03 | 7.5086E-05 | |
| | E | 2.9361E-02 | 3.0610E-03 | <2E-16 | 8.6209E-04 | |
| 4 | PRS_add | 1.6633E-01 | 3.0189E-03 | <2E-16 | 2.7665E-02 | 4.9110E-01 |
| | PRS_gxe | -3.3556E-03 | 3.0144E-03 | 2.6562E-01 | 1.1260E-05 | |
| | PRS_gxe x E | 8.6537E-03 | 2.9707E-03 | 3.5800E-03 | 7.4887E-05 | |
| | E | 2.9396E-02 | 3.0611E-03 | <2E-16 | 8.6413E-04 | |
| 4* | PRS_add | NA | NA | NA | NA | NA |
| | PRS_gxe | NA | NA | NA | NA | |
| | PRS_gxe x E | NA | NA | 2.0000E-03 | NA | |
| | E | NA | NA | NA | NA | |

Table S11: Analysis of WHR/HD

| Model | Component | Estimate | Standard error | P value | R squared | Adjusted R squared |
|---|---|---|---|---|---|---|
| 1 | PRS_trd | 1.6750E-01 | 3.0110E-03 | <2E-16 | 2.8056E-02 | 4.9310E-01 |
| | PRS_trd x E | 3.2810E-03 | 3.2890E-03 | 3.1853E-01 | 1.0765E-05 | |
| | E | -5.4030E-02 | 3.3750E-03 | <2E-16 | 2.9192E-03 | |
| 2 | PRS_add | 1.6600E-01 | 3.0113E-03 | <2E-16 | 2.7556E-02 | 4.9270E-01 |
| | PRS_add x E | 3.2245E-03 | 3.4474E-03 | 3.4962E-01 | 1.0397E-05 | |
| | E | -5.4786E-02 | 3.5299E-03 | <2E-16 | 3.0015E-03 | |
| 3 | PRS_add | 1.6599E-01 | 3.0111E-03 | <2E-16 | 2.7553E-02 | 4.9270E-01 |
| | PRS_gxe x E | 6.8639E-03 | 3.1727E-03 | 3.0510E-02 | 4.7113E-05 | |
| | E | -5.8610E-02 | 3.2629E-03 | <2E-16 | 3.4351E-03 | |
| 4 | PRS_add | 1.6597E-01 | 3.0114E-03 | <2E-16 | 2.7546E-02 | 4.9270E-01 |
| | PRS_gxe | 1.4867E-03 | 3.0110E-03 | 6.2148E-01 | 2.2103E-06 | |
| | PRS_gxe x E | 6.8498E-03 | 3.1728E-03 | 3.0860E-02 | 4.6920E-05 | |
| | E | -5.8604E-02 | 3.2629E-03 | <2E-16 | 3.4344E-03 | |
| 4* | PRS_add | NA | NA | NA | NA | NA |
| | PRS_gxe | NA | NA | NA | NA | |
| | PRS_gxe x E | NA | NA | 2.8000E-02 | NA | |
| | E | NA | NA | NA | NA | |

Table S12: Analysis of WHR/PA

| Model | Component | Estimate | Standard error | P value | R squared | Adjusted R squared |
|---|---|---|---|---|---|---|
| 1 | PRS_trd | 1.4940E-02 | 2.9490E-04 | <2E-16 | 2.2320E-04 | 5.0180E-01 |
| | PRS_trd x E | -3.3440E-04 | 3.2180E-04 | 2.9880E-01 | 1.1182E-07 | |
| | E | -6.3270E-03 | 3.2360E-04 | <2E-16 | 4.0031E-05 | |
| 2 | PRS_add | 1.4320E-02 | 2.9570E-04 | <2E-16 | 2.0506E-04 | 4.9950E-01 |
| | PRS_add x E | -1.6710E-04 | 3.8920E-04 | 6.6763E-01 | 2.7922E-08 | |
| | E | -6.3850E-03 | 3.9060E-04 | <2E-16 | 4.0768E-05 | |
| 3 | PRS_add | 1.4330E-02 | 2.9570E-04 | <2E-16 | 2.0535E-04 | 4.9950E-01 |
| | PRS_gxe x E | 1.7280E-04 | 3.8240E-04 | 6.5130E-01 | 2.9860E-08 | |
| | E | -6.1650E-03 | 3.8390E-04 | <2E-16 | 3.8007E-05 | |
| 4 | PRS_add | 1.4320E-02 | 2.9620E-04 | <2E-16 | 2.0506E-04 | 4.9950E-01 |
| | PRS_gxe | -7.2460E-05 | 2.9450E-04 | 8.0561E-01 | 5.2505E-09 | |
| | PRS_gxe x E | 1.7170E-04 | 3.8240E-04 | 6.5334E-01 | 2.9481E-08 | |
| | E | -6.1650E-03 | 3.8390E-04 | <2E-16 | 3.8007E-05 | |
| 4* | PRS_add | NA | NA | NA | NA | NA |
| | PRS_gxe | NA | NA | NA | NA | |
| | PRS_gxe x E | NA | NA | 6.4900E-01 | NA | |
| | E | NA | NA | NA | NA | |

Table S13: Analysis of CAD/BMI

| Model | Component | Estimate | Standard error | P value | R squared | Pseudo R squared |
|---|---|---|---|---|---|---|
| 1 | PRS_trd | 3.0125E-01 | 1.5478E-02 | <2E-16 | 5.8928E-03 | 1.5557E-01 |
| | PRS_trd x E | -4.4117E-02 | 1.4415E-02 | 2.2100E-03 | 8.9361E-05 | |
| | E | 2.7670E-01 | 1.4352E-02 | <2E-16 | 4.9950E-03 | |
| 2 | PRS_add | 2.9312E-01 | 1.5444E-02 | <2E-16 | 5.5217E-03 | 1.5487E-01 |
| | PRS_add x E | -4.8127E-02 | 1.5068E-02 | 1.4030E-03 | 5.9586E-05 | |
| | E | 2.7609E-01 | 1.4510E-02 | <2E-16 | 5.6526E-03 | |
| 3 | PRS_add | 2.8362E-01 | 1.5017E-02 | <2E-16 | 5.5091E-03 | 1.5479E-01 |
| | PRS_gxe x E | 4.5834E-02 | 1.6357E-02 | 5.0770E-03 | 5.6645E-05 | |
| | E | 3.0985E-01 | 1.6955E-02 | <2E-16 | 5.8821E-03 | |
| 4 | PRS_add | 2.9462E-01 | 1.5401E-02 | <2E-16 | 6.0278E-03 | 1.5515E-01 |
| | PRS_gxe | 5.1311E-02 | 1.5800E-02 | 1.1600E-03 | 2.2946E-04 | |
| | PRS_gxe x E | 3.3844E-02 | 1.6773E-02 | 4.3610E-02 | 6.1422E-05 | |
| | E | 3.0243E-01 | 1.6944E-02 | <2E-16 | 5.9038E-03 | |
| 5 | PRS_add | 2.9399E-01 | 1.5401E-02 | <2E-16 | 6.0267E-03 | 1.5571E-01 |
| | PRS_gxe | 5.0919E-02 | 1.5848E-02 | 1.3130E-03 | 2.2925E-04 | |
| | PRS_gxe x E | 3.0508E-02 | 1.7257E-02 | 7.7082E-02 | 6.0893E-05 | |
| | E | 3.5244E-01 | 2.1415E-02 | <2E-16 | 5.9626E-03 | |
| | E_sq | -6.6539E-02 | 1.7120E-02 | 1.0200E-04 | 5.7760E-07 | |
| 5* | PRS_add | NA | NA | NA | NA | NA |
| | PRS_gxe | NA | NA | NA | NA | |
| | PRS_gxe x E | NA | NA | 7.9000E-02 | NA | |
| | E | NA | NA | NA | NA | |
| | E_sq | NA | NA | NA | NA | |

Table S14: Analysis of CAD/HDL

| Model | Component | Estimate | Standard error | P value | R squared | Pseudo R squared |
|---|---|---|---|---|---|---|
| 1 | PRS_trd | 2.9393E-01 | 1.7495E-02 | <2E-16 | 6.1285E-03 | 1.5971E-01 |
| | PRS_trd x E | 1.1592E-03 | 1.9230E-02 | 9.5193E-01 | 9.5218E-04 | |
| | E | -4.1697E-01 | 2.0216E-02 | <2E-16 | 1.0702E-02 | |
| 2 | PRS_add | 2.6936E-01 | 1.7404E-02 | <2E-16 | 4.9552E-03 | 1.5731E-01 |
| | PRS_add x E | 1.6849E-02 | 2.1356E-02 | 4.3014E-01 | 7.4604E-04 | |
| | E | -4.2647E-01 | 2.1192E-02 | <2E-16 | 1.3039E-02 | |
| 3 | PRS_add | 2.7204E-01 | 1.6113E-02 | <2E-16 | 4.9511E-03 | 1.5790E-01 |
| | PRS_gxe x E | 8.8456E-02 | 2.2329E-02 | 7.4500E-05 | 6.4184E-05 | |
| | E | -3.7969E-01 | 2.3974E-02 | <2E-16 | 9.2092E-03 | |
| 4 | PRS_add | 2.9942E-01 | 1.6834E-02 | <2E-16 | 6.3279E-03 | 1.5920E-01 |
| | PRS_gxe | -1.0409E-01 | 1.8128E-02 | 9.3700E-09 | 9.0783E-04 | |
| | PRS_gxe x E | 3.4037E-02 | 2.4334E-02 | 1.6190E-01 | 6.0183E-05 | |
| | E | -4.1122E-01 | 2.5503E-02 | <2E-16 | 9.2656E-03 | |
| 5 | PRS_add | 2.9935E-01 | 1.6858E-02 | <2E-16 | 6.2032E-03 | 1.6092E-01 |
| | PRS_gxe | -1.0721E-01 | 1.7914E-02 | 2.1700E-09 | 9.3686E-04 | |
| | PRS_gxe x E | 2.9607E-02 | 2.2718E-02 | 1.9248E-01 | 5.2069E-05 | |
| | E | -4.2276E-01 | 2.4075E-02 | <2E-16 | 1.6270E-02 | |
| | E_sq | 1.3892E-01 | 1.8986E-02 | 2.5300E-13 | 3.6606E-03 | |
| 5* | PRS_add | NA | NA | NA | NA | NA |
| | PRS_gxe | NA | NA | NA | NA | |
| | PRS_gxe x E | NA | NA | 1.9900E-01 | NA | |
| | E | NA | NA | NA | NA | |
| | E_sq | NA | NA | NA | NA | |

Table S15: Analysis of CAD/HGB

| Model | Component | Estimate | Standard error | P value | R squared | Pseudo R squared |
|---|---|---|---|---|---|---|
| 1 | PRS_trd | 3.9859E-01 | 1.6547E-02 | <2E-16 | 1.0072E-02 | 9.5891E-02 |
| | PRS_trd x E | -4.9321E-02 | 1.6548E-02 | 2.8790E-03 | 8.2886E-08 | |
| | E | -7.5291E-02 | 1.9754E-02 | 1.3800E-04 | 8.6854E-04 | |
| 2 | PRS_add | 4.0149E-01 | 1.6556E-02 | <2E-16 | 1.0213E-02 | 9.6234E-02 |
| | PRS_add x E | -5.7814E-02 | 1.8494E-02 | 1.7720E-03 | 1.0157E-06 | |
| | E | -9.6483E-02 | 1.9756E-02 | 1.0400E-06 | 9.2229E-04 | |
| 3 | PRS_add | 3.9617E-01 | 1.6443E-02 | <2E-16 | 1.0218E-02 | 9.5942E-02 |
| | PRS_gxe x E | 2.9006E-02 | 1.7991E-02 | 1.0692E-01 | 3.9114E-05 | |
| | E | -9.7965E-02 | 2.0814E-02 | 2.5200E-06 | 1.0666E-03 | |
| 4 | PRS_add | 3.9403E-01 | 1.6518E-02 | <2E-16 | 1.0128E-02 | 9.6016E-02 |
| | PRS_gxe | -2.2148E-02 | 1.6433E-02 | 1.7772E-01 | 2.0122E-05 | |
| | PRS_gxe x E | 3.1856E-02 | 1.8118E-02 | 7.8706E-02 | 3.9269E-05 | |
| | E | -9.8992E-02 | 2.0733E-02 | 1.8000E-06 | 1.0711E-03 | |
| 5 | PRS_add | 3.9420E-01 | 1.6520E-02 | <2E-16 | 1.0126E-02 | 9.7037E-02 |
| | PRS_gxe | -2.1230E-02 | 1.6440E-02 | 1.9656E-01 | 1.9077E-05 | |
| | PRS_gxe x E | 2.6830E-02 | 1.7450E-02 | 1.2401E-01 | 3.5265E-05 | |
| | E | -8.9310E-02 | 2.0050E-02 | 8.3900E-06 | 8.3839E-04 | |
| | E_sq | 7.0910E-02 | 1.4590E-02 | 1.1700E-06 | 5.5147E-04 | |
| 5* | PRS_add | NA | NA | NA | NA | NA |
| | PRS_gxe | NA | NA | NA | NA | |
| | PRS_gxe x E | NA | NA | 1.2100E-01 | NA | |
| | E | NA | NA | NA | NA | |
| | E_sq | NA | NA | NA | NA | |

Table S16: Analysis of CAD/WHR

| Model | Component | Estimate | Standard error | P value | R squared | Pseudo R squared |
|---|---|---|---|---|---|---|
| 1 | PRS_trd | 2.9200E-01 | 1.6693E-02 | <2E-16 | 6.0128E-03 | 1.5448E-01 |
| | PRS_trd x E | 4.4622E-03 | 1.6139E-02 | 7.8200E-01 | 1.6100E-03 | |
| | E | 3.5840E-01 | 1.9946E-02 | <2E-16 | 1.0342E-02 | |
| 2 | PRS_add | 2.6985E-01 | 1.6645E-02 | <2E-16 | 5.0699E-03 | 1.5231E-01 |
| | PRS_add x E | -3.1582E-03 | 1.9285E-02 | 8.6992E-01 | 1.7449E-03 | |
| | E | 3.6778E-01 | 2.1547E-02 | <2E-16 | 1.4775E-02 | |
| 3 | PRS_add | 2.7887E-01 | 1.5184E-02 | <2E-16 | 5.0592E-03 | 1.5286E-01 |
| | PRS_gxe x E | 6.9231E-02 | 1.7245E-02 | 5.9600E-05 | 1.2607E-05 | |
| | E | 3.3266E-01 | 2.1917E-02 | <2E-16 | 9.1268E-03 | |
| 4 | PRS_add | 3.1460E-01 | 1.6240E-02 | <2E-16 | 6.8698E-03 | 1.5423E-01 |
| | PRS_gxe | 1.1150E-01 | 1.7730E-02 | 3.1600E-10 | 9.4661E-04 | |
| | PRS_gxe x E | 1.9690E-02 | 1.8990E-02 | 2.9987E-01 | 1.4105E-05 | |
| | E | 3.5640E-01 | 2.2630E-02 | <2E-16 | 8.9643E-03 | |
| 5 | PRS_add | 3.1396E-01 | 1.6238E-02 | <2E-16 | 6.9290E-03 | 1.5445E-01 |
| | PRS_gxe | 1.1053E-01 | 1.7813E-02 | 5.4700E-10 | 9.6574E-04 | |
| | PRS_gxe x E | 2.0640E-02 | 1.9316E-02 | 2.8530E-01 | 1.6723E-05 | |
| | E | 3.8892E-01 | 2.6283E-02 | <2E-16 | 8.1488E-03 | |
| | E_sq | -4.0173E-02 | 1.5895E-02 | 1.1500E-02 | 1.2382E-03 | |
| 5* | PRS_add | NA | NA | NA | NA | NA |
| | PRS_gxe | NA | NA | NA | NA | |
| | PRS_gxe x E | NA | NA | 3.1300E-01 | NA | |
| | E | NA | NA | NA | NA | |
| | E_sq | NA | NA | NA | NA | |

Table S17: Analysis of DIAB/BMI

| Model | Component | Estimate | Standard error | P value | R squared | Pseudo R squared |
|---|---|---|---|---|---|---|
| 1 | PRS_trd | 3.5821E-01 | 1.9269E-02 | <2E-16 | 5.7302E-03 | 1.9215E-01 |
| | PRS_trd x E | -4.0841E-02 | 1.4630E-02 | 5.2500E-03 | 2.4538E-03 | |
| | E | 7.3823E-01 | 1.4880E-02 | <2E-16 | 4.9856E-02 | |
| 2 | PRS_add | 3.3376E-01 | 1.9104E-02 | <2E-16 | 5.1266E-03 | 1.9026E-01 |
| | PRS_add x E | -4.8023E-02 | 1.8153E-02 | 8.1570E-03 | 1.8207E-03 | |
| | E | 7.3622E-01 | 1.7188E-02 | <2E-16 | 6.4821E-02 | |
| 3 | PRS_add | 3.3378E-01 | 1.7208E-02 | <2E-16 | 5.1080E-03 | 1.9154E-01 |
| | PRS_gxe x E | 1.0459E-01 | 1.6414E-02 | 1.8700E-10 | 1.3064E-04 | |
| | E | 7.0106E-01 | 1.7405E-02 | <2E-16 | 4.9107E-02 | |
| 4 | PRS_add | 3.8949E-01 | 1.8929E-02 | <2E-16 | 7.9691E-03 | 1.9348E-01 |
| | PRS_gxe | 1.4926E-01 | 2.0850E-02 | 8.1300E-13 | 1.4861E-03 | |
| | PRS_gxe x E | 4.6134E-02 | 1.8386E-02 | 1.2100E-02 | 1.2589E-04 | |
| | E | 7.3357E-01 | 1.8431E-02 | <2E-16 | 4.8929E-02 | |
| 5 | PRS_add | 3.8344E-01 | 1.8903E-02 | <2E-16 | 8.1591E-03 | 1.9652E-01 |
| | PRS_gxe | 1.4593E-01 | 2.1251E-02 | 6.5600E-12 | 1.5792E-03 | |
| | PRS_gxe x E | 3.9114E-02 | 1.8740E-02 | 3.6872E-02 | 1.7740E-04 | |
| | E | 9.0371E-01 | 2.7123E-02 | <2E-16 | 3.3058E-02 | |
| | E_sq | -1.4769E-01 | 1.6514E-02 | <2E-16 | 5.2314E-03 | |
| 5* | PRS_add | NA | NA | NA | NA | NA |
| | PRS_gxe | NA | NA | NA | NA | |
| | PRS_gxe x E | NA | NA | 5.7000E-02 | NA | |
| | E | NA | NA | NA | NA | |
| | E_sq | NA | NA | NA | NA | |

Table S18: Analysis of DIAB/HDL

| Model | Component | Estimate | Standard error | P value | R squared | Pseudo R squared |
|---|---|---|---|---|---|---|
| 1 | PRS_trd | 3.4762E-01 | 2.1746E-02 | <2E-16 | 8.0258E-03 | 1.5315E-01 |
| | PRS_trd x E | -2.6718E-02 | 2.3479E-02 | 2.5515E-01 | 3.5421E-03 | |
| | E | -8.2062E-01 | 2.5082E-02 | <2E-16 | 2.7943E-02 | |
| 2 | PRS_add | 3.1218E-01 | 2.1655E-02 | <2E-16 | 5.5667E-03 | 1.4774E-01 |
| | PRS_add x E | 1.6912E-02 | 2.3801E-02 | 4.7737E-01 | 1.6835E-03 | |
| | E | -8.5768E-01 | 2.4851E-02 | <2E-16 | 3.1046E-02 | |
| 3 | PRS_add | 3.2930E-01 | 1.8212E-02 | <2E-16 | 5.6055E-03 | 1.4913E-01 |
| | PRS_gxe x E | 1.2084E-01 | 2.1389E-02 | 1.6100E-08 | 2.2715E-05 | |
| | E | -8.1255E-01 | 2.6048E-02 | <2E-16 | 2.6700E-02 | |
| 4 | PRS_add | 3.8444E-01 | 1.9600E-02 | <2E-16 | 8.8963E-03 | 1.5185E-01 |
| | PRS_gxe | -1.8018E-01 | 2.3111E-02 | 6.3700E-15 | 2.0603E-03 | |
| | PRS_gxe x E | 1.1440E-02 | 2.5764E-02 | 6.5703E-01 | 2.0703E-05 | |
| | E | -8.5320E-01 | 2.7699E-02 | <2E-16 | 2.6667E-02 | |
| 5 | PRS_add | 3.8678E-01 | 1.9687E-02 | <2E-16 | 8.7198E-03 | 1.5654E-01 |
| | PRS_gxe | -1.8274E-01 | 2.2256E-02 | <2E-16 | 2.0285E-03 | |
| | PRS_gxe x E | 9.7332E-03 | 2.2961E-02 | 6.7164E-01 | 2.6457E-05 | |
| | E | -8.1068E-01 | 2.4824E-02 | <2E-16 | 4.8685E-02 | |
| | E_sq | 2.6556E-01 | 2.0868E-02 | <2E-16 | 1.2719E-02 | |
| 5* | PRS_add | NA | NA | NA | NA | NA |
| | PRS_gxe | NA | NA | NA | NA | |
| | PRS_gxe x E | NA | NA | 6.7300E-01 | NA | |
| | E | NA | NA | NA | NA | |
| | E_sq | NA | NA | NA | NA | |

Table S19: Analysis of DIAB/HGB

| Model | Component | Estimate | Standard error | P value | R squared | Pseudo R squared |
|---|---|---|---|---|---|---|
| 1 | PRS_trd | 3.9859E-01 | 1.6547E-02 | <2E-16 | 1.0072E-02 | 9.5891E-02 |
| | PRS_trd x E | -4.9321E-02 | 1.6548E-02 | 2.8790E-03 | 8.2886E-08 | |
| | E | -7.5291E-02 | 1.9754E-02 | 1.3800E-04 | 8.6854E-04 | |
| 2 | PRS_add | 4.0149E-01 | 1.6556E-02 | <2E-16 | 1.0213E-02 | 9.6234E-02 |
| | PRS_add x E | -5.7814E-02 | 1.8494E-02 | 1.7720E-03 | 1.0157E-06 | |
| | E | -9.6483E-02 | 1.9756E-02 | 1.0400E-06 | 9.2229E-04 | |
| 3 | PRS_add | 3.9617E-01 | 1.6443E-02 | <2E-16 | 1.0218E-02 | 9.5942E-02 |
| | PRS_gxe x E | 2.9006E-02 | 1.7991E-02 | 1.0692E-01 | 3.9114E-05 | |
| | E | -9.7965E-02 | 2.0814E-02 | 2.5200E-06 | 1.0666E-03 | |
| 4 | PRS_add | 3.9403E-01 | 1.6518E-02 | <2E-16 | 1.0128E-02 | 9.6016E-02 |
| | PRS_gxe | -2.2148E-02 | 1.6433E-02 | 1.7772E-01 | 2.0122E-05 | |
| | PRS_gxe x E | 3.1856E-02 | 1.8118E-02 | 7.8706E-02 | 3.9269E-05 | |
| | E | -9.8992E-02 | 2.0733E-02 | 1.8000E-06 | 1.0711E-03 | |
| 5 | PRS_add | 3.9420E-01 | 1.6520E-02 | <2E-16 | 1.0126E-02 | 9.7037E-02 |
| | PRS_gxe | -2.1230E-02 | 1.6440E-02 | 1.9656E-01 | 1.9077E-05 | |
| | PRS_gxe x E | 2.6830E-02 | 1.7450E-02 | 1.2401E-01 | 3.5265E-05 | |
| | E | -8.9310E-02 | 2.0050E-02 | 8.3900E-06 | 8.3839E-04 | |
| | E_sq | 7.0910E-02 | 1.4590E-02 | 1.1700E-06 | 5.5147E-04 | |
| 5* | PRS_add | NA | NA | NA | NA | NA |
| | PRS_gxe | NA | NA | NA | NA | |
| | PRS_gxe x E | NA | NA | 1.4000E-01 | NA | |
| | E | NA | NA | NA | NA | |
| | E_sq | NA | NA | NA | NA | |

Table S20: Analysis of DIAB/WHR

| Model | Component | Estimate | Standard error | P value | R squared | Pseudo R squared |
|---|---|---|---|---|---|---|
| 1 | PRS_trd | 3.5821E-01 | 2.0268E-02 | <2E-16 | 6.6562E-03 | 1.7495E-01 |
| | PRS_trd x E | -1.3966E-02 | 1.7942E-02 | 4.3634E-01 | 3.8136E-03 | |
| | E | 9.8241E-01 | 2.2310E-02 | <2E-16 | 6.9247E-02 | |
| 2 | PRS_add | 2.8932E-01 | 2.0128E-02 | <2E-16 | 4.5401E-03 | 1.6951E-01 |
| | PRS_add x E | -3.9748E-03 | 1.9380E-02 | 8.3749E-01 | 2.6245E-03 | |
| | E | 1.0050E+00 | 2.2644E-02 | <2E-16 | 8.1054E-02 | |
| 3 | PRS_add | 3.0808E-01 | 1.7317E-02 | <2E-16 | 4.5603E-03 | 1.7019E-01 |
| | PRS_gxe x E | 6.6845E-02 | 1.5879E-02 | 2.5600E-05 | 1.3638E-05 | |
| | E | 9.9016E-01 | 2.2454E-02 | <2E-16 | 7.0119E-02 | |
| 4 | PRS_add | 3.8260E-01 | 1.9380E-02 | <2E-16 | 8.1314E-03 | 1.7309E-01 |
| | PRS_gxe | 1.9480E-01 | 2.2410E-02 | <2E-16 | 1.9210E-03 | |
| | PRS_gxe x E | -1.6120E-02 | 1.8580E-02 | 3.8550E-01 | 1.4095E-05 | |
| | E | 1.0080E+00 | 2.2890E-02 | <2E-16 | 6.9265E-02 | |
| 5 | PRS_add | 3.7980E-01 | 1.9360E-02 | <2E-16 | 8.3553E-03 | 1.7367E-01 |
| | PRS_gxe | 1.9300E-01 | 2.2660E-02 | <2E-16 | 1.9664E-03 | |
| | PRS_gxe x E | -1.5010E-02 | 1.8960E-02 | 4.2847E-01 | 2.7283E-05 | |
| | E | 1.0810E+00 | 3.0210E-02 | <2E-16 | 6.3901E-02 | |
| | E_sq | -6.9300E-02 | 1.7740E-02 | 9.3700E-05 | 8.1614E-03 | |
| 5* | PRS_add | NA | NA | NA | NA | NA |
| | PRS_gxe | NA | NA | NA | NA | |
| | PRS_gxe x E | NA | NA | 4.4200E-01 | NA | |
| | E | NA | NA | NA | NA | |
| | E_sq | NA | NA | NA | NA | |

Table S21: Analysis of HYP/BMI

| Model | Component | Estimate | Standard error | P value | R squared | Pseudo R squared |
|---|---|---|---|---|---|---|
| 1 | PRS_trd | 3.0435E-01 | 1.0387E-02 | <2E-16 | 1.2837E-02 | 2.3239E-01 |
| | PRS_trd x E | -2.9421E-02 | 1.3202E-02 | 2.5847E-02 | 2.6229E-04 | |
| | E | 5.4732E-01 | 1.3607E-02 | <2E-16 | 3.8648E-02 | |
| 2 | PRS_add | 3.2212E-01 | 1.0372E-02 | <2E-16 | 1.4400E-02 | 2.3478E-01 |
| | PRS_add x E | -2.1023E-02 | 1.2021E-02 | 8.0320E-02 | 2.3812E-04 | |
| | E | 5.6348E-01 | 1.2420E-02 | <2E-16 | 4.2978E-02 | |
| 3 | PRS_add | 3.1980E-01 | 1.0252E-02 | <2E-16 | 1.4370E-02 | 2.3477E-01 |
| | PRS_gxe x E | 1.7057E-02 | 1.0166E-02 | 9.3378E-02 | 1.1595E-05 | |
| | E | 5.5393E-01 | 1.0470E-02 | <2E-16 | 4.6545E-02 | |
| 4 | PRS_add | 3.2775E-01 | 1.0403E-02 | <2E-16 | 1.5113E-02 | 2.3521E-01 |
| | PRS_gxe | 4.8037E-02 | 1.0397E-02 | 3.8300E-06 | 3.3492E-04 | |
| | PRS_gxe x E | 9.9565E-03 | 1.0283E-02 | 3.3293E-01 | 1.2541E-05 | |
| | E | 5.5324E-01 | 1.0461E-02 | <2E-16 | 4.6561E-02 | |
| 5 | PRS_add | 3.2761E-01 | 1.0406E-02 | <2E-16 | 1.5110E-02 | 2.3594E-01 |
| | PRS_gxe | 4.8018E-02 | 1.0421E-02 | 4.0700E-06 | 3.3476E-04 | |
| | PRS_gxe x E | 8.8240E-03 | 1.0226E-02 | 3.8818E-01 | 1.2310E-05 | |
| | E | 5.9917E-01 | 1.3063E-02 | <2E-16 | 4.7011E-02 | |
| | E_sq | -7.3970E-02 | 1.2184E-02 | 1.2700E-09 | 3.9390E-06 | |
| 5* | PRS_add | NA | NA | NA | NA | NA |
| | PRS_gxe | NA | NA | NA | NA | |
| | PRS_gxe x E | NA | NA | 4.0100E-01 | NA | |
| | E | NA | NA | NA | NA | |
| | E_sq | NA | NA | NA | NA | |

Table S22: Analysis of HYP/HDL

| Model | Component | Estimate | Standard error | P value | R squared | Pseudo R squared |
|---|---|---|---|---|---|---|
| 1 | PRS_trd | 3.1889E-01 | 1.0833E-02 | <2E-16 | 1.5775E-02 | 1.8968E-01 |
| | PRS_trd x E | -2.4890E-04 | 1.4012E-02 | 9.8583E-01 | 3.3782E-04 | |
| | E | -3.3435E-01 | 1.5594E-02 | <2E-16 | 1.3948E-02 | |
| 2 | PRS_add | 3.1683E-01 | 1.0818E-02 | <2E-16 | 1.5440E-02 | 1.8933E-01 |
| | PRS_add x E | 1.1904E-02 | 1.4769E-02 | 4.2025E-01 | 2.1948E-04 | |
| | E | -3.5863E-01 | 1.6425E-02 | <2E-16 | 1.5671E-02 | |
| 3 | PRS_add | 3.1581E-01 | 1.0736E-02 | <2E-16 | 1.5425E-02 | 1.8953E-01 |
| | PRS_gxe x E | 3.9938E-02 | 1.3690E-02 | 3.5300E-03 | 2.1322E-04 | |
| | E | -3.2691E-01 | 1.4611E-02 | <2E-16 | 1.5909E-02 | |
| 4 | PRS_add | 3.1619E-01 | 1.0740E-02 | <2E-16 | 1.5465E-02 | 1.8958E-01 |
| | PRS_gxe | -1.5519E-02 | 1.0697E-02 | 1.4684E-01 | 5.8220E-05 | |
| | PRS_gxe x E | 3.7207E-02 | 1.3818E-02 | 7.0900E-03 | 2.1104E-04 | |
| | E | -3.2839E-01 | 1.4688E-02 | <2E-16 | 1.5926E-02 | |
| 5 | PRS_add | 3.1654E-01 | 1.0762E-02 | <2E-16 | 1.5321E-02 | 1.9298E-01 |
| | PRS_gxe | -1.7129E-02 | 1.0693E-02 | 1.0918E-01 | 6.3936E-05 | |
| | PRS_gxe x E | 3.2533E-02 | 1.3307E-02 | 1.4490E-02 | 1.8052E-04 | |
| | E | -3.8647E-01 | 1.5157E-02 | <2E-16 | 2.5452E-02 | |
| | E_sq | 1.4590E-01 | 1.1784E-02 | <2E-16 | 4.0924E-03 | |
| 5* | PRS_add | NA | NA | NA | NA | NA |
| | PRS_gxe | NA | NA | NA | NA | |
| | PRS_gxe x E | NA | NA | 1.6000E-02 | NA | |
| | E | NA | NA | NA | NA | |
| | E_sq | NA | NA | NA | NA | |

Table S23: Analysis of HYP/HGB

| Model | Component | Estimate | Standard error | P value | R squared | Pseudo R squared |
|---|---|---|---|---|---|---|
| 1 | PRS_trd | 3.3074E-01 | 1.0162E-02 | <2E-16 | 1.7192E-02 | 1.7439E-01 |
| | PRS_trd x E | -2.7941E-02 | 1.2712E-02 | 2.7956E-02 | 1.8961E-08 | |
| | E | 5.7026E-02 | 1.5155E-02 | 1.6800E-04 | 9.7330E-05 | |
| 2 | PRS_add | 3.2936E-01 | 1.0159E-02 | <2E-16 | 1.7047E-02 | 1.7420E-01 |
| | PRS_add x E | -2.7562E-02 | 1.2515E-02 | 2.7643E-02 | 8.0486E-08 | |
| | E | 5.7767E-02 | 1.4962E-02 | 1.1300E-04 | 1.1063E-04 | |
| 3 | PRS_add | 3.2767E-01 | 1.0125E-02 | <2E-16 | 1.7048E-02 | 1.7410E-01 |
| | PRS_gxe x E | 7.4835E-03 | 1.0396E-02 | 4.7160E-01 | 8.0821E-06 | |
| | E | 4.1394E-02 | 1.2712E-02 | 1.1290E-03 | 1.2370E-04 | |
| 4 | PRS_add | 3.2741E-01 | 1.0132E-02 | <2E-16 | 1.7026E-02 | 1.7411E-01 |
| | PRS_gxe | -6.8117E-03 | 9.9709E-03 | 4.9451E-01 | 4.4791E-06 | |
| | PRS_gxe x E | 7.8976E-03 | 1.0414E-02 | 4.4822E-01 | 7.9976E-06 | |
| | E | 4.1487E-02 | 1.2717E-02 | 1.1060E-03 | 1.2340E-04 | |
| 5 | PRS_add | 3.2743E-01 | 1.0136E-02 | <2E-16 | 1.6992E-02 | 1.7500E-01 |
| | PRS_gxe | -6.8641E-03 | 9.9727E-03 | 4.9127E-01 | 4.5463E-06 | |
| | PRS_gxe x E | 6.2777E-03 | 1.0278E-02 | 5.4135E-01 | 5.0805E-06 | |
| | E | 4.4917E-02 | 1.2577E-02 | 3.5500E-04 | 2.3403E-04 | |
| | E_sq | 6.3697E-02 | 1.0227E-02 | 4.7200E-10 | 7.9044E-04 | |
| 5* | PRS_add | NA | NA | NA | NA | NA |
| | PRS_gxe | NA | NA | NA | NA | |
| | PRS_gxe x E | NA | NA | 5.3500E-01 | NA | |
| | E | NA | NA | NA | NA | |
| | E_sq | NA | NA | NA | NA | |

Table S24: Analysis of HYP/WHR

| Model | Component | Estimate | Standard error | P value | R squared | Pseudo R squared |
|---|---|---|---|---|---|---|
| 1 | PRS_trd | 3.1752E-01 | 1.0380E-02 | <2E-16 | 1.4492E-02 | 2.1475E-01 |
| | PRS_trd x E | -1.3876E-02 | 1.3165E-02 | 2.9191E-01 | 6.7864E-04 | |
| | E | 6.0347E-01 | 1.6420E-02 | <2E-16 | 4.6703E-02 | |
| 2 | PRS_add | 3.1087E-01 | 1.0345E-02 | <2E-16 | 1.3818E-02 | 2.1387E-01 |
| | PRS_add x E | -2.2827E-02 | 1.2898E-02 | 7.6751E-02 | 4.2083E-04 | |
| | E | 6.2666E-01 | 1.6190E-02 | <2E-16 | 5.1526E-02 | |
| 3 | PRS_add | 3.0917E-01 | 1.0150E-02 | <2E-16 | 1.3839E-02 | 2.1418E-01 |
| | PRS_gxe x E | 4.5484E-02 | 1.0729E-02 | 2.2400E-05 | 1.4691E-04 | |
| | E | 5.9877E-01 | 1.4158E-02 | <2E-16 | 5.5254E-02 | |
| 4 | PRS_add | 3.1793E-01 | 1.0350E-02 | <2E-16 | 1.4765E-02 | 2.1459E-01 |
| | PRS_gxe | 4.5702E-02 | 1.0359E-02 | 1.0300E-05 | 3.9083E-04 | |
| | PRS_gxe x E | 3.6072E-02 | 1.0939E-02 | 9.7500E-04 | 1.4508E-04 | |
| | E | 6.0063E-01 | 1.4199E-02 | <2E-16 | 5.5086E-02 | |
| 5 | PRS_add | 3.1790E-01 | 1.0350E-02 | <2E-16 | 1.4773E-02 | 2.1460E-01 |
| | PRS_gxe | 4.5609E-02 | 1.0363E-02 | 1.0800E-05 | 3.9575E-04 | |
| | PRS_gxe x E | 3.6173E-02 | 1.0948E-02 | 9.5300E-04 | 1.3722E-04 | |
| | E | 6.0270E-01 | 1.4722E-02 | <2E-16 | 5.3586E-02 | |
| | E_sq | -5.9040E-03 | 1.0974E-02 | 5.9055E-01 | 7.7262E-04 | |
| 5* | PRS_add | NA | NA | NA | NA | NA |
| | PRS_gxe | NA | NA | NA | NA | |
| | PRS_gxe x E | NA | NA | 1.1200E-03 | NA | |
| | E | NA | NA | NA | NA | |
| | E_sq | NA | NA | NA | NA | |
